# Centuries of genome instability and evolution in soft-shell clam transmissible cancer

**DOI:** 10.1101/2022.08.07.503107

**Authors:** Samuel F.M. Hart, Marisa A. Yonemitsu, Rachael M. Giersch, Brian F. Beal, Gloria Arriagada, Brian W. Davis, Elaine A. Ostrander, Stephen P. Goff, Michael J. Metzger

## Abstract

Transmissible cancers are infectious parasitic clones of malignant cells that metastasize to new hosts, living past the death of the founder animal in which the cancer initiated. Several lineages of transmissible cancer have recently been identified in bivalves, including one that has spread through the soft-shell clam (*Mya arenaria*) population along the east coast of North America. To investigate the evolutionary history of this transmissible cancer lineage, we assembled a highly contiguous 1.2 Gb soft-shell clam reference genome and characterized somatic mutations from cancer sequences. We show that all cancer cases observed descend from a single founder and cluster into two geographically distinct sub-lineages. We discover a previously unreported clock-like mutational signature that predicts the cancer lineage to be 344 to 877 years old, indicating that it spread undetected long before it was first observed in the 1970s. We observe high mutation density, widespread copy number gain, structural rearrangement, loss of heterozygosity, variable telomere lengths, mitochondrial genome expansion, and transposable element activity, all indicative of an unstable cancer genome. Our study reveals the ability for an invertebrate cancer lineage to survive for centuries while its genome continues to structurally mutate, likely contributing to the ability of this lineage to adapt as a parasitic cancer.

**SUMMARY:** The genome of a contagious cancer in clams reveals structural instability of multiple types throughout the ∼500 years since its origin.

## INTRODUCTION

Most cancers arise from oncogenic mutations in host cells and remain confined to the body of that host. However, a small number of transmissible cancer lineages exist in which cancer cells metastasize repeatedly to new hosts and live past the death of their original hosts (*1*). Observed cases of transmissible cancer in nature include Canine Transmissible Venereal Tumor (CTVT) in dogs (*2*), two unrelated lineages of Devil Facial Tumor Disease (DFTD) in Tasmanian devils (*3, 4*), and at least eight Bivalve Transmissible Neoplasia (BTN) lineages observed in several marine bivalve species (*5–9*). Although transmissible cancers and their host genomes have been well characterized in dogs (*10–12*) and devils (*13–15*), little is known about the evolutionary history of the BTN lineages, which have only recently been recognized as transmissible cancers. Here we perform the first genome-wide analysis of a BTN lineage by focusing on the single lineage found in the soft-shell clam (*<u>M</u>ya <u>ar</u>enaria*), or MarBTN.

BTN is a fatal leukemia-like cancer characterized by the proliferation of cancer cells in the circulatory fluid of the bivalve and dissemination into tissues in the later stages of disease. BTN cells can survive for days to weeks in seawater (*16, 17*) and they likely spread from animal to animal by transmission through the water column. This cancer, referred to in the literature as disseminated neoplasia or hemic neoplasia, was first reported in soft-shell clams in the 1970s (*18, 19*) and has since been found across much of the soft-shell clam native range along the east coast of North America (**Fig. 1A**). All cancer isolates tested in a 2015 study were shown to be of clonal origin, and it has been hypothesized that historical observations of the cancer dating back to the 1970s were occurrences of this same clonal lineage (*5*). However, it is not known how long this lineage has propagated, or how the genome has evolved since the original cancer initiated. To address these and other questions, we assembled a high-quality soft-shell clam reference genome and characterized the genome evolution of the MarBTN lineage by comparative analysis of healthy clam and MarBTN sequences. We show a striking pattern of mutation occurrence and evolution, suggestive of an unstable genome with the potential to rapidly mutate.

**Figure 1.**
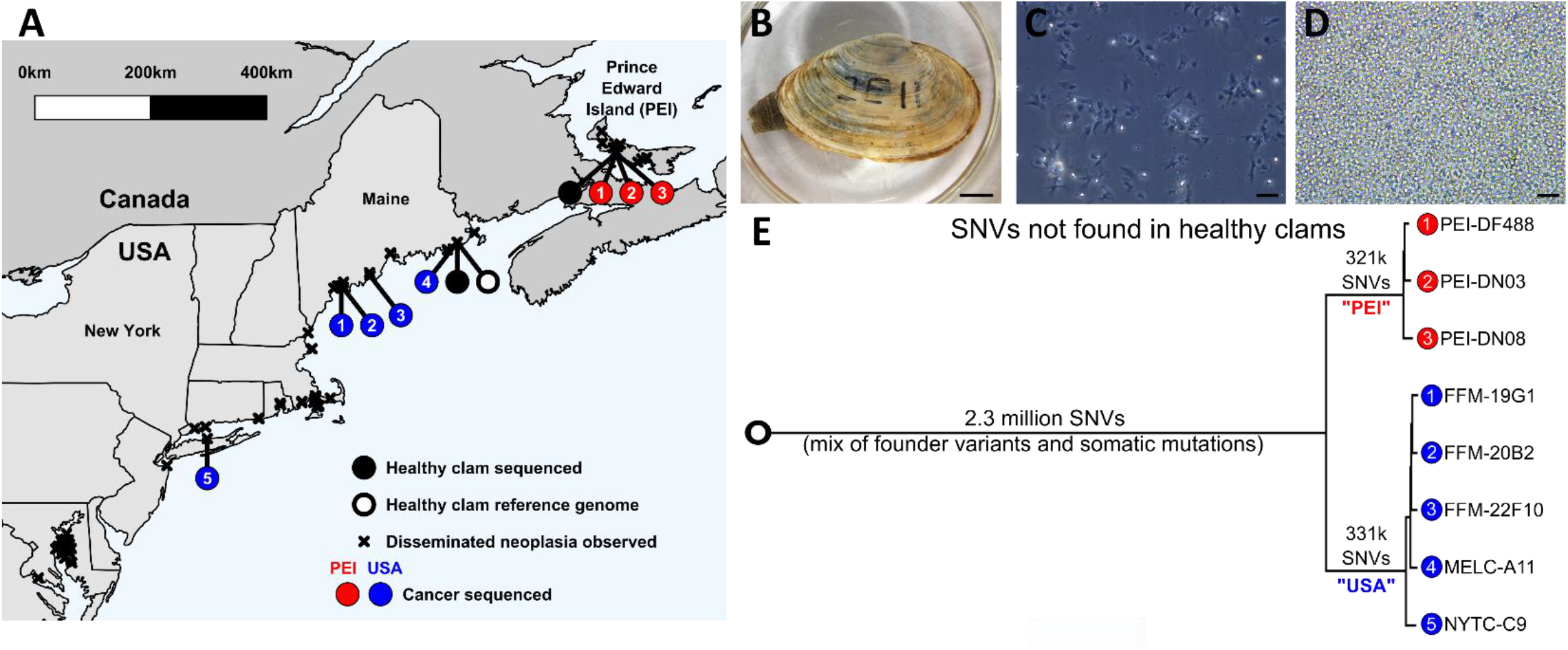
MarBTN distribution and sequencing. **(A)** Locations of samples sequenced (circles) and disseminated neoplasia observations (x’s) along the east coast of North America. Circles colored for healthy clams (black) and MarBTN sampled from the PEI (red) or USA (blue) coast. **(B)** Image of healthy clam sequenced and used to assemble reference genome (MELC-2E11) and **(C)** hemolymph of the same clam, with hemocytes extending pseudopodia. **(D)** Hemolymph from a clam infected with MarBTN (FFM-22F10), with distinct rounded morphology and lack of pseudopodia of cancer cells. Scale bars are 10 mm for the clam and 50 µm for the hemolymph. **(E)** Phylogeny of cancer samples built from pairwise differences of SNVs not found in healthy clams, excluding regions that show evidence of LOH. Numbers along branches indicate the number of SNVs unique to and shared by individuals in that clade.

## RESULTS

### Sample sequencing and genome assembly

We assembled a soft-shell clam reference genome from a single healthy female clam collected from Larrabee Cove, Machiasport, Maine, USA (**Fig. 1B-C**, MELC-2E11). We assembled PacBio long reads into contigs using FALCON-Unzip (*20*), scaffolded contigs to the chromosome-level with Hi-C sequences using FALCON-Phase (**Supplementary Fig. 1**), polished the scaffolds using 10X Chromium reads, and annotated with RNAseq reads using Maker to yield a high quality reference genome. The final reference genome is 1.22 Gb organized into 17 phased scaffolds, matching the 17 chromosomes expected based on karyotype data (*21*). The contig N50 is 3.4 Mb and the metazoan BUSCO (Benchmarking Universal Single Copy Orthologs (*22*)) score is 94.9%. Our assembly is similar in size, GC, and repeat content of a recently published *Mya arenaria* genome (*23*) but with drastically improved contiguity and completeness (**Supplementary Table 1**), allowing for comprehensive genomic investigation into the evolutionary history of MarBTN.

We performed whole genome sequencing (WGS) on three healthy uninfected clams and eight isolates of MarBTN from the hemolymph of highly infected clams (e.g. **Fig. 1D**) from five locations across the established MarBTN range (*24*) (**Fig. 1A, Supplementary Table 2**), and called single nucleotide variants (SNVs) against the reference genome. Contaminating host variants were removed from MarBTN sequences via variant calling thresholds, rather than using paired tissue sequences as has been done for other transmissible cancers, since MarBTN hemolymph isolates were highly pure (>97% cancer DNA) while paired tissue samples from the host often contained high amounts of cancer DNA due to dissemination (up to 72% cancer DNA) (**Supplementary Fig. 2**).

To investigate somatic evolution of the MarBTN lineage, it is important to distinguish between founder variants, those present in the genome of the founder clam from which the cancer initially arose, and somatic mutations, which occurred during the propagation and evolution of the cancer lineage. 10.7 million SNVs were shared by all MarBTN samples but not present in the reference genome. Of these, 8.1 million were found in at least one of the three healthy clams, indicating that these variants are likely founder variants.

A MarBTN phylogeny, built from pairwise SNV differences between samples, confirmed the previous analysis identifying two distinct sub-lineages of MarBTN (*5*), here referred to as the Prince Edward Island (PEI) and United States of America (USA) sub-lineages **(Fig. 1E)**. Most known founder SNVs (those also found in healthy animals) were present in both sub-lineages of MarBTN, but we observed some genomic regions with clusters of founder SNVs in one sub-lineage but not the other. These are unlikely to be somatic mutations, instead they likely indicate loss of heterozygosity (LOH) events which took place after divergence of the sub-lineages. LOH was identified in 8% and 13% of the USA and PEI sub-lineage genomes **(Supplementary Fig. 3)**, respectively, and these regions were excluded during identification of somatic mutations in the following SNV analysis. Remaining SNVs found in all cancer samples, but no healthy clams, represent a mix of both founder variants and somatic mutations (2.3 million), while SNVs found in just one or the other sub-lineage represent likely somatic mutations (700 k). The majority of these SNVs were shared by all individuals in a sub-lineage and are herein referred to as “high confidence somatic mutations” (321 k for PEI and 331 k for USA).

### Mutational biases in MarBTN

We observed a distinct SNV mutational bias in somatic mutations within both the PEI and USA sub-lineages that was not found in healthy clams (**Fig. 2B)**. These biases are nearly identical between the two sub-lineages and were also present in more recent mutations, such as SNVs unique to each MarBTN sample **(Supplementary Fig. 4)**. *De novo* signature extraction, which deconvolutes mutational biases in their trinucleotide context between samples (*25*), yielded four mutational signatures (**Supplementary Fig. 5)**. Three signatures were found in both healthy clams and MarBTN samples, and thus are likely endogenous within the germline of clam genomes. One signature closely resembles COSMIC signature 1 (termed Sig1’), with characteristic bias for C>T mutations at CpG sites and associated with the deamination of methylated CpGs in humans (*26*). Sig1’ represents a greater fraction of mutations in the PEI sub-lineage **(Supplementary Fig. 6)**, which may indicate that PEI has more methylated CpG sites than USA. Sig1’ also represents a greater fraction of mutations in coding regions **(Supplementary Fig. 7)**, fitting prior observations that methylation is elevated in gene regions in bivalves (*27*). The other two signatures are “flatter” and less distinctive, somewhat resembling COSMIC signatures 5 and 40, which are both associated with aging in humans (*28, 29*).

**Figure 2.**
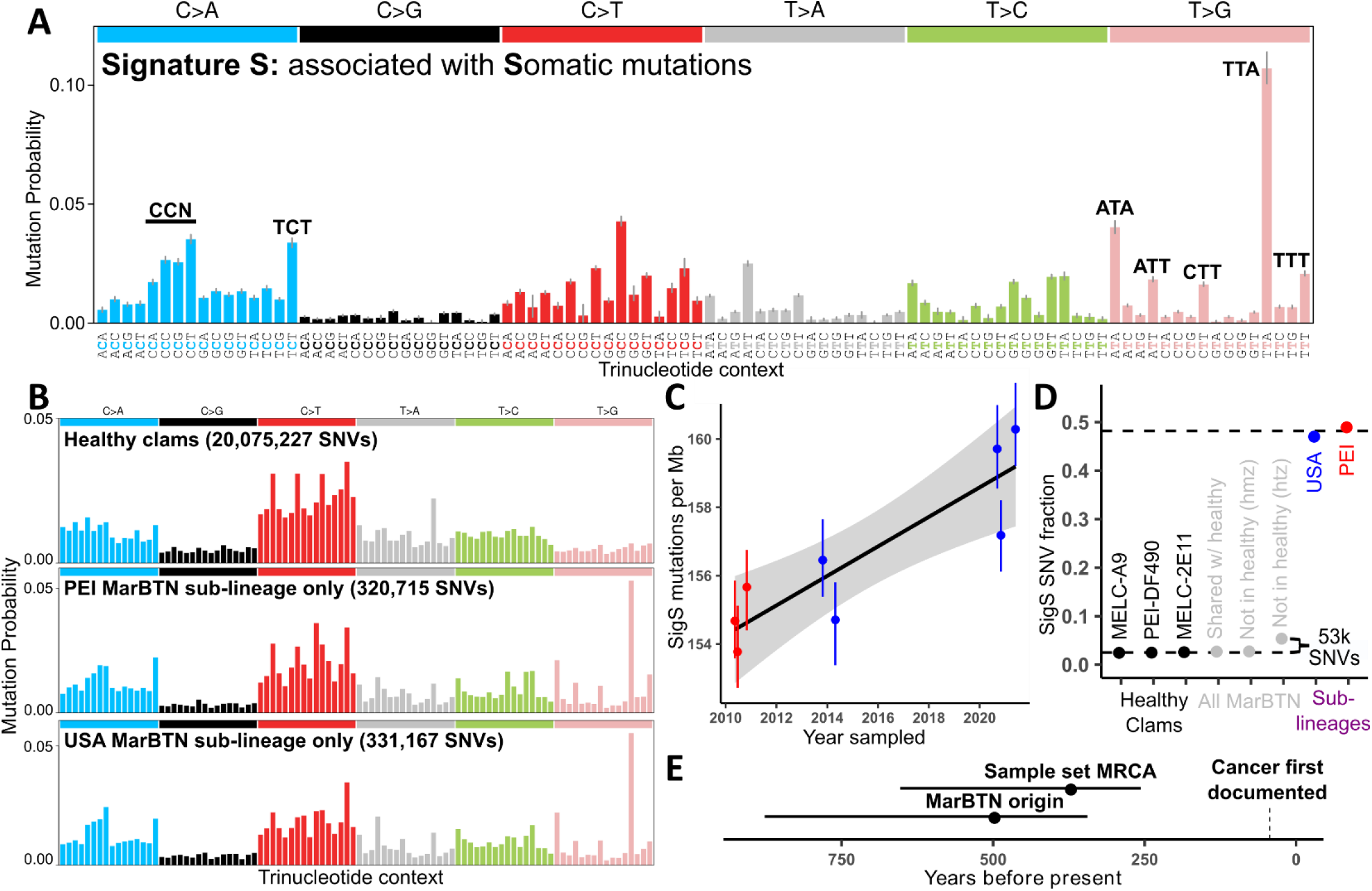
Unique mutational signature found in somatic mutations dates cancer to ∼500 years old. **(A)** *De novo* extracted mutational biases for signature S. **(B)** Trinucleotide context of SNVs found in healthy clams (top) and high confidence somatic mutations in PEI (middle) or USA (bottom) sub-lineages, corrected for mutational opportunities in the clam genome. Trinucleotide order same as in A. **(C)** Signature S mutations per Mb across MarBTN samples correlated with sampling date, with linear regression and 95% confidence interval (grey) overlaid. SNVs found in healthy clams, all MarBTN samples, or LOH regions are excluded. **(D)** Fraction of SNVs attributed to signature S from healthy clams (black), variants found in all MarBTN samples (grey), and high confidence somatic mutations (colored). Variants found in all MarBTN samples are divided by whether they are found in healthy clams and whether they are homozygous (hmz) or heterozygous (htz). Dashed lines display signature S fraction estimates for likely somatic mutations and likely founder variants. **(E)** Age estimate of the most recent common ancestor (MRCA) of the USA and PEI sub-lineages and of BTN origin from signature S mutations. Error bars in all plots display 95% confidence intervals.

A single signature captured the biases specific to the <u>S</u>omatic mutations in MarBTN, termed SigS (**Fig. 2A)**. The closest analog in the COSMIC database of human mutational signatures is signature 9, which shares a T>G bias in A/T trinucleotide contexts (*28*). Signature 9 SNVs in humans represents mutations induced by polymerase eta during somatic hypermutation (*28*). This may indicate that an error-prone polymerase with similar biases to human polymerase eta is involved in the cancer’s replication. Polymerase eta is also involved in translesion synthesis in humans (*30*), so SigS could alternatively represent mutations introduced by error-prone repair of DNA lesions during MarBTN replication. In addition to the striking T>G bias in A/T contexts, there is also a notable bias towards C>A mutations compared to healthy clam SNVs, particularly CC>CA and TCT>TAT. Interestingly, both C>A and T>G mutations have been linked to oxidative DNA damage (*31*). Clam hemolymph is strongly hypoxic in late stages of the disease (*32*), so this environment may also be contributing to these mutational biases.

### MarBTN is several centuries old

Previous studies have used signature 1 mutation accumulation, which is considered clock-like in humans and other mammals (*33, 34*), to date CTVT origin to 4,000-8,500 years before present (*11*). However, Sig1’ mutations in MarBTN have not accumulated in a clock-like manner (**Supplementary Fig. 8**). This may be due to methylation changes affecting CpG>TpG mutation rates and/or inherent differences between clams and mammals. However, SigS-attributed mutations since the divergence of the two sub-lineages correlate with sampling date, accumulating at a rate of 432 (+/- 188) mutations/Gb/year over the 11 years separating our sample set (**Fig. 2C**). This corresponds to the sub-lineages diverging 369 years ago (95% CI: 257-653 years), long before the first recorded observations of disseminated neoplasia in soft-shell clams in the 1970s (*18, 19*).

We also estimated how long prior to the sub-lineage divergence the cancer first arose in the founder clam and began horizontal transmission. SigS contributed roughly half of high confidence somatic mutations in each sub-lineage but was virtually absent from SNVs in the healthy clam population (**Fig. 2D)**. If we assume the SigS mutation rate has remained constant since oncogenesis and that the founder clam SNVs have a similar profile of genomic SNVs to those observed in healthy clams, we can estimate that 3.1% of heterozygous SNVs found in all cancer samples, but no healthy clams, are somatic mutations attributed to SigS. This corresponds to 126 years by the rate estimate above, for a total cancer age estimate of 495 years (95% CI: 344-877 years) (**Fig. 2E)**.

If we also assume the fraction of SigS somatic mutations has remained constant at 47% since oncogenesis, we estimate that 6.8% (95% CI: 6.5-7.1%) of all heterozygous SNVs found in all cancer samples, but no healthy clams, are somatic mutations, for a total somatic SNV estimate of 441 and 452 mu/Mb for the PEI and USA sub-lineages, respectively. This is a much higher mutation density than that estimated for the <40-year-old DFTD lineages (DFT1: <3.1 mu/Mb, DFT2: <1.3 mu/Mb) (*14*), but less than the >4000-year-old CTVT (∼867 mu/Mb from exome data) (*11*), showing that mutation density generally scales with age across the small number of characterized transmissible cancer lineages.

### Selection on SNVs is largely neutral

We used the ratio of non-synonymous to synonymous coding changes (dN/dS) to infer selection acting on coding regions in our sample set. After correcting for mutational opportunities in coding regions, a ratio of one indicates neutral selection, >1 indicates positive selection, and <1 indicates negative/purifying selection. We used dNdScv(*35*) to determine that the global dN/dS for healthy clam SNVs was 0.454 (95% CI: 0.451-0.457), indicating genes are generally under negative selection in clam genomes, as expected. On a gene-by-gene basis, 70% of intact coding genes (16,222/23,273) in healthy clams have significantly negative dN/dS, while 0.4% (88/23,273) are significantly positive. Genes under positive selection in hosts may indicate genes at the host-pathogen interface that are under selection for continued novelty. In the case of clams, some of these genes may be a response to MarBTN evolution itself, though this hypothesis cannot be tested by the current study.

High confidence somatic mutations had a global dN/dS of 0.982 (95% CI: 0.943-1.024), indicating that MarBTN is largely dominated by neutral selection, reflecting observations in human cancers (*36*) and CTVT(*11*) **(Supplementary Fig. 9)**. We found no genes with a dN/dS ratio significantly <1, indicating no genes under significant negative, or purifying, selection, but we did identify five genes with a dN/dS ratio significantly >1, indicating positive selection (**Supplementary Table 3**). For all five of these genes, nearly all somatic mutations were found on a single haplotype in a single sub-lineage. None of these genes are under positive selection in healthy clams, suggesting that these are truly under positive selection in only a single sub-lineage and not founder or host clam SNVs. A particularly notable gene among the five is a TEN1-like gene under positive selection in just the USA sub-lineage. TEN1 is a component of the CTC1-STN1-TEN1 complex, which plays a crucial role in telomere replication and genome stability (*37*).

### Widespread structural mutation

Polyploidy has been described in disseminated neoplasia in several bivalve species (*24, 38*). In *Mya arenaria*, disseminated neoplasia cells have approximately double the chromosome count and genome content of healthy clam cells (*21*). Given the discovery that these cells are of clonal origin (*5*), we had hypothesized that a full genome duplication occurred early in the cancer’s evolution and that most of the MarBTN genome should be 4N. To test this theory, we called copy number states across each non-reference sample genome based on read depth **(Fig. 3A)**. As expected, both healthy clams were 2N across nearly the entire genome **(Fig. 3B)**. Unexpectedly, MarBTN samples displayed a wide variety of copy number states.

**Figure 3.**
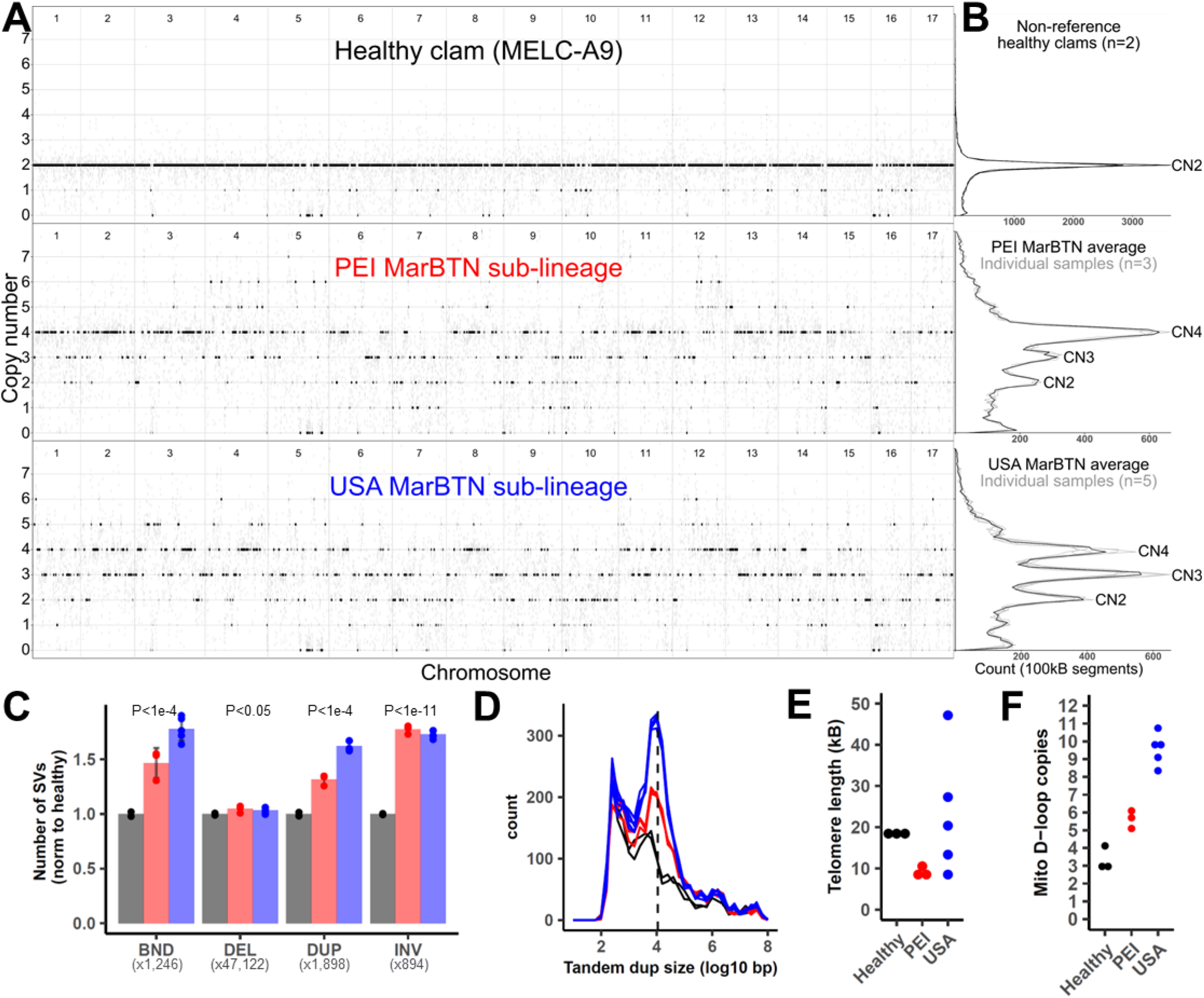
Widespread copy number gain and structural mutation. **(A)** Copy number calls across clam genome, rounded to the nearest integer (black) and unrounded (grey) in 100 kB segments. The healthy clam is a representative individual, and the MarBTN sub-lineages are averages of each individual sample from that sub-lineage, which were in close agreement. **(B)** Summary of copy number states across entire genomes for two non-reference healthy clams and MarBTN sub-lineages. Grey lines display copy number summaries for individual samples within each sub-lineage, which are in close agreement. **(C)** Number of called SVs in each sample after removing SVs found in the reference clam. Values are normalized to the average number of SVs in non-reference healthy clams for each SV type (numbers below SV type labels). P-values are from unequal variance t-test between BTN samples (n=8) and non-reference healthy clams (n=2). Labels follow delly abbreviations of SV types: BND = translocations, DEL = deletions, DUP = tandem duplications, INV = Inversions. Error bars indicate standard deviation. **(D)** Size distribution of tandem duplications in each non-reference sample. Dashed line indicates 11 kB. **(E)** Telomere length estimated by TelSeq for each sample. **(F)** Tandem duplicate copies of the mitochondrial D-loop region per sample. Healthy normal clams are black, MarBTN from PEI are red, and MarBTN samples from USA are blue. Samples are ordered in same order as supplementary table 2.

PEI samples were predominantly 4N with substantial 3N and 2N portions, while USA samples were more evenly distributed between 4N, 3N, and 2N **(Fig. 3B)**. Copy number calls in cancer samples displayed close agreement within sub-lineages (R^2^>0.94), but they revealed large differences between sub-lineages, despite still positively correlating (R^2^=0.53-0.56) **(Supplementary Fig. 10)**. Variant allele frequencies (VAF) for high confidence somatic mutations largely support copy number calls **(Supplementary Fig. 11)**, with some off-target VAF peaks, most notably in the lower copy number regions (<3N) indicating some of these regions are higher copy number than called, likely due to lower read mapping in polymorphic genome regions.

Many mid-chromosome breakpoints were apparent in the copy number calls, indicating that the MarBTN genome has likely undergone widespread structural alterations in addition to whole-chromosome and within-chromosome copy number gain. We are unable to resolve the structure of the MarBTN genome with the short reads of our current data set but were able to call likely structural variants (SVs) from split reads. Relative to non-reference healthy clams, MarBTN samples had a significantly higher number of deletions, inversions, tandem duplications, and inter-chromosomal translocations, indicating substantial somatic structural alterations **(Fig. 3C)**.

Comparing likely somatic structural variants specific to each sub-lineage, USA samples had significantly more translocations and tandem duplications than PEI **(Supplementary Fig. 12)**. Median somatic tandem duplication sizes displayed a distinct distribution around a mode of ∼11 kB **(Fig. 3D, Supplementary Fig. 13)**. In human cancers, tandem duplication phenotypes of this same size distribution are thought to be driven by the loss of TP53 and BRCA1 (*39*), indicating that a parallel mutational process may be influencing the observed genome instability in MarBTN and more active in the USA sub-lineage.

Maintenance of telomere length is a requirement for an immortalized cell line such as MarBTN and would be necessary for long-term survival. We estimated telomere lengths for each sample and found them to be highly variable within the USA sub-lineage (8-47 kB), while short but relatively stable within the PEI sub-lineage (8-11 kB) in comparison to healthy clams (18-19 kB) **(Fig. 3E)**. Variable telomere lengths in the USA sub-lineage may be mediated by the TEN1-like gene that is under positive selection in that sub-lineage, as the CTC1-STN1-TEN1 complex inhibits telomerase and is involved in telomere length homeostasis (*37*).

### Mitochondrial genome evolution

A tree built from pairwise mitochondrial SNV differences between samples reflects a similar phylogeny to that built from genomic SNVs **(Supplementary Fig. 16)**. This indicates no evidence of mitochondrial uptake or recombination with host mitochondria, which has been observed in other transmissible cancers (*7, 40, 41*). Transitions were highly overrepresented in both healthy and cancer samples, with C>T mutations comprising 41/50 likely somatic mutations **(Supplementary Fig. 17)**. Somatic mutations resulted in missense mutations in at least 10 of the 12 mitochondrial genes and appear to be under relaxed selection, with dN/dS ratios of 0.97 (95%CI: 0.45-2.1) versus 0.26 (95%CI: 0.11-0.58) for SNVs in healthy clams **(Supplementary Fig. 18)**.

When aligned to the published *Mya arenaria* mitochondrial genome (*42*), short read sequences from all MarBTN and healthy samples display an increase in coverage across the mitochondrial D-loop (**Supplementary Fig. 19)**, indicating the region is multi-copy. The D-loop is part of the non-coding control region of the mitochondrial genome and is the origin of both replication and transcription. We resolved this region with PacBio long reads from the healthy reference clam, revealing three copies in tandem. Two of the D-loop copies contain a 236 bp insertion not found in the published mitochondrial genome. The insert includes an 80 bp region with 70% guanine content, likely complicating previous PCR-based efforts to resolve it. Altogether, the observed copies extend the D-loop region of the reference clam genome from 845 bp to 2,727 bp and the full mitochondrial genome to 19,815 bp.

Read coverage of the D-loop region suggest that there have been additional somatic tandem duplications in the MarBTN mitogenome. While read coverage indicates three or four copies in the non-reference healthy clams, PEI MarBTN samples have 5-6 copies and USA MarBTN samples have 8-11 (**Fig. 3F)**. These somatic tandem duplications likely arose via replication errors and the trend towards increased copies in cancer suggests that they may be under selection. Selection can act on the level of the mitogenome itself, giving it a replicative advantage over other mitogenomes (as hypothesized for CTVT), or on the level of the cancer cell, if this duplication provides cancer cells a replicative advantage over others. Notably, the suspected mitogenome site under selection during repeated mitochondrial capture in CTVT is hypothesized to also be a control region polymorphism (*41*).

### Transposable element mobilization

MarBTN is known to contain the LTR-retrotransposon *Steamer* at a much higher copy number than healthy clams, indicating likely somatic expansion (*43*). To test whether *Steamer* activity is ongoing we identified *Steamer* insertion sites using split reads spanning *Steamer* and the reference genome. Only 5-11 sites were found in healthy samples, versus 275-460 sites in cancer samples. One hundred ninety-three sites are shared by all cancer samples, indicating that *Steamer* expansion likely began early in the cancer’s evolution, while sub-lineage-specific *Steamer* integrations indicate that *Steamer* has continued to replicate somatically in the MarBTN genome (**Fig. 4A**). However, *Steamer* has generated more insertions within the USA sub-lineage (248) than the PEI sub-lineage (64), indicating the regulatory environments of the sub-lineages have not remained stable since they diverged.

**Figure 4.**
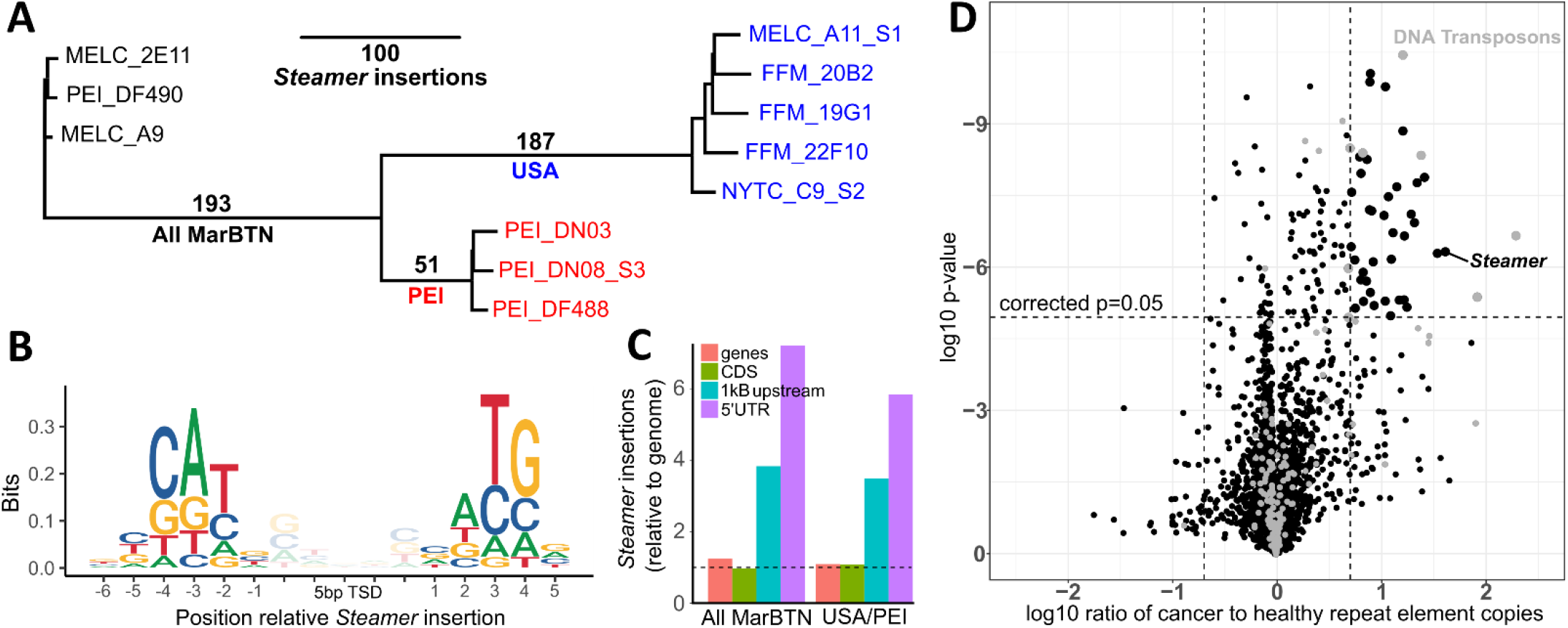
Somatic expansions of *Steamer* and other Tes. **(A)** Phylogeny of all samples built from pairwise differences of *Steamer* insertion sites, colored by healthy (black), USA MarBTN (blue), and PEI MarBTN (red). Numbers along branches indicate the number of insertions unique to and shared by individuals in that clade. **(B)** Nucleotide relative to the 5 bp target site duplication (TSD) of all *Steamer* insertions. Probabilities are normalized by nucleotide content of the genome. **(C)** *Steamer* insertion probability in annotated genome regions, normalized by read mapping rates and relative to full genome. Displayed for insertions found in all MarBTN samples but no healthy clams, and unique to each sub-lineage but shared by all individual in that sub-lineage. Dashed line indicates expectation given random insertions. **(D)** Volcano plot comparing copy number of all repeat elements in MarBTN and healthy clam samples. Dashed lines correspond to significance threshold (p=0.05, Bonferroni corrected) and 5-fold differences. Elements annotated as DNA transposons are marked in grey.

We also observed strong biases for *Steamer* to insert at specific genomic sequences. *Steamer* has a palindromic bias for NATG outside the 5 bp target site duplication (CATNnnnnnNATG), inserting at these locations 45× more frequently than expected by chance (**Fig. 4B)**. *Steamer* was also >3× more likely to insert in the 1000 bp upstream of genes than would be expected by chance (**Fig. 4C)**. We also observed early *Steamer* insertions (those found in all MarBTN samples) upstream of cancer-associated orthologs more often than expected by chance in the reverse but not the forward orientation **(Supplementary Fig. 14)**. This bias that could indicate either an insertion preference for those locations or a selective advantage to MarBTN cells with those insertions.

We further investigated whether other transposable elements (TEs) in addition to *Steamer* have expanded somatically by identifying a library of TEs found in clam genomes and counting the copy number of each TE type in each sample. Forty-five TEs were present at a significantly higher copy number in cancer samples relative to healthy clams, after removing TEs with less than five-fold differences (**Fig. 4D)**. TEs annotated as DNA transposons were enriched in this data set (8/45: 17.8%) compared to the total TE library (171/4471: 3.8%), indicating this TE type may have been particularly successful in somatically expanding its copy number in MarBTN. LTR retrotransposons (like *Steamer*) appear to have had more success in the USA versus PEI sub-lineage. Thirty-six TEs have significantly more copies in the USA sub-lineage than PEI, and eight of those are LTR-retrotransposons, compared to zero LTR-retrotransposons out of 20 of those more highly expanded in PEI **(Supplementary Fig. 15)**. Lower copy numbers of LTR retrotransposons and other TEs in the PEI sub-lineage could be linked to the increased methylation indicated by mutational signature analysis, as methylation is thought to repress TE mobilization (*27, 44*). Our finding of widespread increases in TE copy numbers across sub-lineages complements our structural mutation results, both indicating genome instability of the MarBTN lineage and more mutations of certain types in the USA sub-lineage.

## DISCUSSION

Our genome analyses reveal a diverse set of somatic mutations occurring in MarBTN, with continued accumulation of SNVs and widespread structural mutations indicative of genome instability. Prior research has implicated p53 homolog mis-regulation as a possible driver of this cancer (*45*), and our work supports this hypothesis. We find signatures in the MarBTN genome similar to signatures associated with p53 loss in humans; tolerance of widespread copy number gain (*46*), TE expansions (*47*), and ∼11 kB tandem duplications (*39*). However, it is unclear whether these mutations have consistently occurred over time or have been generated in multiple punctuated chromothripsis-like events. Genomic studies of the dog and devil transmissible cancers have observed contrastingly stable genomes, remaining predominantly diploid even despite thousands of years of somatic evolution in the case of CTVT (*10, 48*). Polyploidy has been reported in other BTN lineages in other bivalves (*24*), indicating that genome instability may be a common driver mechanism or a tolerated by-product of conserved processes in BTN evolution.

Interestingly, we observe differences in the amount of structural mutations, telomere length, and TE amplification between the two sub-lineages, indicating genome instability or mutation tolerance may have changed over time in MarBTN after the sub-lineages diverged. This may be in part mediated by the TEN1-like gene under positive selection in the USA sub-lineage. These changes in fundamental mutational mechanisms in different sub-lineages post-divergence highlight the fact that oncogenesis is not a single event, but an ongoing evolutionary process.

In contrast to the above unstable and variable structural processes, we observe a pattern of consistent single nucleotide mutation biases in both sub-lineages. Most notable in these biases is the novel profile and clock-like nature of mutational signature S. We hypothesize that this signature is due in part to an error-prone polymerase involved in MarBTN replication (due to its similarity to human COSMIC signature 9 caused by polymerase eta). Its clock-like quality may be due to consistent MarBTN replication rates over time, as seen in human somatic cells with defective proofreading polymerases (*49*) or due to continual damage of chromosomal DNA and its repair using translesion synthesis. Using SigS, we estimate that this cancer lineage is 344-877 years old, and was present well before it was first observed in the 1970s (*18, 19*). This places MarBTN at an intermediate age compared with DFTD (<40 years (*15*)) and CTVT (4000-8500 years (*11*)).

We observe that the MarBTN genome is largely dominated by neutral selection, reflecting observations in human cancers (*36*) and CTVT (*11*), with a few notable genes under positive selection in a single sub-lineage, which may reflect selection for repeated mutations involved in critical oncogenic processes. However, we also note that selection is not simply relevant at the level of the cancer cell, but also on the level of the gene (as seen in MarBTN TE expansions), mitochondria (as seen in CTVT horizontal transfer (*41*) and now MarBTN mitogenome expansion), and hosts (as seen in DFTD (*50*)). Further analysis of MarBTN and other cancers will help us to understand how these selective forces interact to influence cancer evolution, and perhaps how we can manipulate those forces to our advantage in the fight against conventional and transmissible cancers.

Our analysis of the MarBTN genome is presented alongside an independent analysis of two lineages in the common cockle (*Cerastoderma edule*), or CedBTN, in a companion paper (*51*). The CedBTN genomes display signatures of ongoing instability similar to what we observe in MarBTN, supporting the hypothesis that genome instability is a common feature of BTN evolution and confirming that long-term survival of a cancer lineage can be maintained despite remarkably widespread and continued genome rearrangement. This level of instability might be expected to lead to an error catastrophe (*52*), yet these cancers have continued to survive and grow for centuries, changing our understanding of what is possible in cancer evolution. Tandem duplications in the mitochondrial control region were also observed in both studies and may represent convergent evolution driven by similar selective mechanisms. Similar tandem duplications in the D-loop have also been observed in human cancers (*53, 54*), and the functional consequences of these mutations remains unclear. Repeated expansion of this region in independent BTN lineages, along with their long history of co-evolution with their hosts, make BTNs unique model systems for the understanding of the functional significance of mitogenome mutations on cancer cell growth and the potential for selfish selection at the level of the mitogenome in cancer.

In contrast, our finding of a specific clock-like mutational signature likely due to an error-prone polymerase, the repeated mutations in a TEN1 homolog likely leading to variable telomere length, and amplification of the *Steamer* retrotransposon and other TEs may be unique features of the BTN in clams. Given the apparent abundance of BTNs, continuing to analyze BTN lineages in other species may reveal both common and unique pathways that have allowed these cancers to repeatedly circumvent new host immune systems and spread through host populations as contagious cancers. These cancers therefore provide unique models for the understanding of cancer evolution and examples of what genomic changes are possible in long-lived cancers evolving together with their hosts.

## ACKNOWLEDGEMENTS

We thank Carol Reinisch and James Sherry for sample collection, Charles Walker for advice and aid during the initiation of this project, Phase Genomics (Shawn Sullivan, Emily Reister, Kyle Langford, and Hayley Mandelson) for HiC scaffolding, Andrew Banman and Nikita Sakhanenko for in-house computing support at PNRI, Sam White and Steven Roberts for consultation on the use of MAKER for gene annotation, Kelley Harris and Jed Carlson for consultation on mutational signatures, and Adrian Baez-Ortega for consultation on somatypus and sigfit.

## AUTHOR CONTRIBUTIONS

S.F.M.H, M.J.M. and S.P.G. contributed to study conceptualization and design. M.J.M., B.F.B., G.A., and S.P.G. contributed to sample collection. M.A.Y. performed high molecular weight extractions. B.W.D, E.A.O., and M.J.M. contributed to 10X sequencing and analysis. M.J.M. assembled the reference genome. R.M.G. and S.F.M.H. contributed to disseminated neoplasia literature search for map. S.F.M.H. performed the genomic analysis. S.F.M.H. wrote the original draft of the manuscript. All authors (S.F.M.H, M.A.Y., R.M.G, B.F.B., G.A., B.W.D., E.A.O., S.P.G., and M.J.M.) contributed to review and editing of the manuscript.

## DATA AVAILABILITY

All code and data not included in the main text or the supplementary materials is available on github (https://github.com/sfhart33/MarBTNgenome) or NCBI (Bioproject accession number forthcoming).

## SUPPLEMENTARY MATERIALS

Materials and Methods

Supplemental Figures 1-22

Supplemental Tables 1-4

References (*55-105*)

## MATERIALS AND METHODS

### Data availability

All code is available on GitHub (https://github.com/sfhart33/MarBTNgenome), including all dependencies with version numbers. Individual commands for genome assembly are listed below with triangular bullets, and scripts corresponding to written genome analysis methods are listed in bullets at the end of each written section. Analysis was performed with an on-premises Linux server running Ubuntu 16.04. The Linux server was equipped with four Intel Xeon Gold 6148 CPUs and 250 GiB system memory.

Raw sequence data is available on the sequence read archive (accession numbers forthcoming). The assembled genome, gene annotations, transcriptome, updated mitogenome, and TE library are available in an NCBI repository (accession numbers forthcoming). Other important data outputs (e.g vcf file of all SNVs, copy number calls, LOH calls, SV calls, *Steamer* insertion sites…) can be obtained by running the supplied code on the raw data or on request. Note that code was written for our institutes working environment and thus some scripts may need to be altered manually to reproduce this analysis.

### *Mya arenaria* genome assembly

#### Reference animal collection and HMW DNA extraction

The clam chosen as the reference animal (MELC-2E11, 62 mm shell length, **Fig. 1B**) was collected from Larrabee Cove, Machiasport, Maine, USA in June 2018, and shipped to the Pacific Northwest Research Institute labs. Hemolymph was drawn from the animal from the pericardial sinus using a 0.5 in 26 gauge needle on a 3 ml syringe, and it was checked for the presence of MarBTN through morphological analysis (**Fig. 1C**) and with a highly sensitive cancer-specific qPCR assay (described in (*16*)). There was no evidence of detectible BTN through either method. Examination of gonad region revealed the presence of eggs, showing that this individual was female.

High molecular weight (HMW) DNA was extracted from ∼50 mg snap-frozen mantle tissue using a modified CTAB extraction protocol (adapted from (*55*)). CTAB isolation buffer (2% CTAB, 1.4 M NaCl, 20 mM EDTA, 100 mM Tris-HCl, pH 8.0) was preheated to 60°C in a water bath. Tissue was minced, then ground with a pestle in 500 µL 60°C CTAB isolation buffer in a 1.7 ml microcentrifuge tube. 20 µL proteinase K was added and the sample was incubated at 60°C for 10 h on a shaker (200 rpm), then held at room temperature. Sample was extracted once with the addition of 500 µL chloroform-isoamyl alcohol (24:1), mixing gently but thoroughly. This produces two phases, an upper aqueous phase which contains the DNA, and a lower chloroform phase that contains some degraded proteins, lipids, and many secondary compounds. The sample was spun at 6,000 × g for 10 min at room temperature to concentrate phases. Aqueous phase was removed with a wide bore pipet, transferred to a new microcentrifuge tube. 2/3 volumes cold isopropanol (237 µL) was added and inverted gently to precipitate nucleic acids. HMW DNA was spooled out with a glass hook and transferred to a 2 mL microcentrifuge tube containing 1 mL wash buffer (76% ethanol, 10 mM ammonium acetate) for 20 minutes. HMW DNA was spun down (6,000 × g for 10 min) after a minimum of 20 min of washing. Supernatant was poured off carefully and allowed to air dry briefly at room temperature. HMW DNA was resuspended in 200 µL TE (10 mM Tris-HCl, 1 mM EDTA, pH 8.0). RNase A (DNase-free, 10 mg/mL, Thermo Scientific, Waltham, MA) was added to a final concentration of 25 ug/ml (0.5µL) and incubated 30 min at 37°C. Sample was diluted to 2 volumes with TE, 10 M ammonium acetate was added to a final concentration of 2.5 M, sample was mixed, 1.2 mL 100% ethanol was added, and sample was gently inverted to precipitate HMW DNA. HMW was spun down (10,000 × g for 10 min at 4°C). Sample was air dried and resuspended in 200 µL TE buffer overnight at 4°C.

#### 10X Chromium sequencing and unsuccessful assembly

High molecular weight genomic DNA (gDNA) was isolated from reference animal (MELC-2E11) tissue using the MagAttract HMW DNA Kit (Qiagen), quantified using Qubit 2.0 (Life Technologies) and fragment size determined using the Agilent 2200 TapeStation. Average fragment size exceeded 50 Kb. Approximately 1 ng of DNA was loaded on the Chromium Genome Chip (10X Genomics). Whole genome sequencing libraries were prepared using Chromium Genome Library & Gel Bead Kit v.2, Chromium i7 Multiplex Kit and Chromium Controller according to 10X Genomics instructions. The resulting library was indexed and sequenced on 0.75 lanes of a single flow cell on the Illumina HiSeq X Ten system, generating 150-bp paired-end reads.

An attempt to assemble a genome of the same reference animal using 10X sequencing was unsuccessful at creating a highly contiguous genome. However, we report it here for transparency of our assembly attempts. *De novo* assembly was performed using Supernova (v2.1.1). Assembly was conducted with the command:

➢ supernova run --id=10X_MELC-2E11-100 -- fastqs=/home/mmetzger/10XChromiumSized/Data --description=“MELC-2E11 sized 10X assembly “ --maxreads=all --accept-extreme-coverage

Secondary attempts at assembly were conducted using down-sampled subsets of the data (75%, 666928259 reads; 50%, 444618839 reads, and 25%, 222309414 reads), using the read numbers listed above for the option “--maxreads”. The pseudohap2 output was used (Mar.3.1.1 Myaare100B_pseudohap2.1.fasta).

#### FALCON-Unzip diploid assembly

HMW DNA extracted using the CTAB protocol was sequenced using the PacBio core facility at the University of Washington Department of Genome Sciences. Subreads were converted to fasta using:

➢ bam2fasta -o Marenaria.3.2 *.subreads.bam

and the FALCON-Unzip pipeline was run to generate a diploid-aware *de novo* assembly using the commands:

➢ fc_run fc_run_marenaria.cfg &> run1.log &
➢ mv all.log all0.log
➢ fc_unzip.py fc_unzip_marenaria.cfg &> run1.std &

Several modifications were made to default configuration parameters, including changing to “pwatcher_type=blocking” and lowering the memory per job in fc_unzip_marenaria.cfg. Configuration files are found on github (fc_run_marenaria.cfg and fc_unzip_marenaria.cfg). FALCON-Unzip version was pbbioconda-0.0.5 and was used with python 3.7. This FALCON-Unzip pipeline resulted in a primary contig assembly (Mar.3.2.2_cns_p_ctg.fasta) and an alternate haplotig assembly (Mar.3.2.2_cns_h_ctg.fasta).

High heterozygosity in species such as bivalves can lead to under-calling of haplotype homology. The purge_haplotigs pipeline(*56*) was used to remove pairs of contigs that were called as separate primary contigs by FALCON-Unzip but which are more likely to be alternate alleles. This tool identifies pairs of syntenic contigs and moves one from the primary assembly to the haplotig assembly generating a new curated assembly (Mar.3.2.3_curated.FALC.fasta).

#### Scaffolding with Hi-C using FALCON-Phase

Chromatin conformation capture data was generated using a Phase Genomics (Seattle, WA) Proximo Hi-C Animal Kit, which is a commercially available version of the Hi-C protocol(*57*). Following the manufacturer’s instructions, intact cells from two adductor muscle samples from the reference clam (MELC-2E11) were crosslinked using a formaldehyde solution, digested using the Sau3AI restriction enzyme, and proximity ligated with biotinylated nucleotides to create chimeric molecules composed of fragments from different regions of the genome that were physically proximal *in vivo*, but not necessarily genomically proximal. Molecules were pulled down with streptavidin beads and processed into an Illumina-compatible sequencing library. Sequencing was performed on an Illumina NextSeq 500, generating a total of 313,340,002 PE150 read pairs.

The Hi-C reads, primary contigs, and alternate haplotigs (Mar.3.2.3_curated.haplotigs.FALC.fasta) were provided as input to FALCON-Phase (https://phasegenomics.github.io/2019/09/19/hic-alignment-and-qc.html) to correct likely phase switching errors. All other options were set to default, except for the options which specify restriction enzyme motifs in the library (GATC) and the number of iterations to perform (100,000,000). Phased contigs were output in pseudohap format, creating one complete set of contigs for each phase.

Reads were aligned to the resulting Phase 0 contig assembly metzger_mussel.phased.0.fasta following Phase Genomics’ standard Hi-C alignment protocol (*58*). Briefly, reads were aligned using BWA-MEM(*59*) with the -5SP and -t 8 options specified, and all other options default. SAMBLASTER (*60*) was used to flag PCR duplicates, which were later excluded from analysis. Alignments were then filtered with samtools (*61*) using the -F 2304 filtering flag to remove non-primary and secondary alignments. These alignments, along with the primary contigs and alternate haplotigs were used as inputs to the scaffolding process.

Phase Genomics’ Proximo Hi-C genome scaffolding platform was used to create chromosome-scale scaffolds from the Phase 0 assembly, following the same single-phase scaffolding procedure described in Bickhart et al. (*62*). As in the LACHESIS method (*63*), this process computes a contact frequency matrix from the aligned Hi-C read pairs, normalized by the number of restriction sites (GATC) on each contig, and constructs scaffolds in such a way as to optimize expected contact frequency and other statistical patterns in Hi-C data. Approximately 120,000 separate Proximo runs were performed to optimize chromosome assignment and scaffold construction in order to make the scaffolds as concordant with the observed Hi-C data as possible. This process resulted in a set of 17 chromosome-scale scaffolds containing 1,212 Mbp of sequence (99.89% of the phase 0 assembly) with a scaffold N50 of 78.4 Mbp. Juicebox (*64, 65*) was used to correct likely scaffolding errors, though no breaks for mis-joined contigs were introduced at this stage in order to maintain exact contig relationships with the Phase 1 assembly.

Separately, Hi-C data were aligned to a concatenated Phase 0 and Phase 1 assembly using the standard protocol cited above. Because this would cause Hi-C data for most homozygous regions to have a MAPQ of 0 (among possible other issues), this alignment emphasizes phase-specific Hi-C relationships. These alignments and the Phase 0 scaffolds were passed to FALCON-Phase’s bamfilt (-f 20 -m 10), bam2 binmat (default options), and phase (-n 100000000 -s 10) steps to generate new phasing metadata intended to correct latent phasing issues not detected during the earlier contig phasing step.

Juicebox was again used to correct remaining scaffolding errors in Phase 0, including introducing a single break into each of eight suspected mis-joined contigs and two breaks into one suspected double-misjoined contig, based on the appearance of Hi-C signals consistent with chimeric joins. These scaffolding changes were replicated to Phase 1, and new scaffolds for each phase were generated using the juicebox_assembly_converter.py script (https://github.com/phasegenomics/juicebox_scripts). In these final scaffolds, both Phase 0 and Phase 1 included 17 scaffolds spanning 99.7% (1,204 Mbp in Phase 0 and 1,214 Mbp in Phase 1) of input with a scaffold N50 of 70.2 Mbp in Phase 0 and 71.4 Mbp in Phase 1 (Mar.3.3.2_p0_PGA_assembly.fasta and Mar.3.3.2_p1_PGA_assembly.fasta). The 17 scaffolds from both of these two haploid assemblies were compiled into Mar.3.3.2_p0p1_PGA_assembly_17.fasta using a custom perl script two_fasta_prefix_compile_firstX.pl

➢ perl two_fasta_prefix_compile_firstX.pl Mar.3.3.2_p0_PGA_assembly.fasta Mar.3.3.2_p1_PGA_assembly.fasta p0_ p1_ Mar.3.3.2_p0p1_PGA_assembly_17.fasta 17

#### Genome Gap-Filling and Polishing using long read PacBio data and 10X linked-read data

PBJelly was run to gap-fill the scaffolded assembly using pbsuite (*66*) (v15.8.24, slightly modified: https://github.com/esrice/PBJelly) using blasr (v5.1) and networkx (v2.2) with Python 2.7, with the protocol file Protocol_MELC.xml. Only captured gaps were filled (no inter-scaffold gaps) using the option “--capturedOnly” during the “support” step. PBJelly was run with the commands:

➢ Jelly.py setup Protocol_MELC.xml
➢ Jelly.py mapping Protocol_MELC.xml
➢ Jelly.py support Protocol_MELC.xml -x “--capturedOnly”
➢ Jelly.py extraction Protocol_MELC.xml
➢ Jelly.py assembly Protocol_MELC.xml -x “--nproc=20”
➢ Jelly.py output Protocol_MELC.xml

The output of PBJelly (Mar.3.3.3_jelly.out.fasta) renamed all scaffolds to Contig0-Contig33, so names were corrected manually based on PBJelly liftOverTable.json (Mar.3.3.3_jelly.out_name.fasta), and the two haploid genomes were separated (using commands listed in PBJellyRenaming.txt) to generate the haploid gap-filled assemblies (Mar.3.3.3.p0.fasta and Mar.3.3.3.p1.fasta)

Direct use of short reads to polish a highly heterozygous genome is likely to introduce more errors than it corrects, due to the mapping of reads from both haplotypes to a single haploid genome. Therefore, we used a phase-aware polishing strategy, using the 10X linked reads generate above, modified from the pipeline described in the vertebrate genome project (https://github.com/VGP/vgp-assembly/tree/master/pipeline/freebayes-polish). Both phases (p0 and p1) of the scaffolded, and gap-filled diploid genome were concatenated into a single diploid reference file with 34 scaffolds, and the linked-read-aware mapper Longranger (*67*) (v2.2.2) was used to map the 10X reads to the diploid gap-filled assembly (Mar.3.3.3_jelly.out_name.fasta), and the output was indexed using samtools (v1.9). FreeBayes (v1.3.1) and Bcftools (v1.10.2) were used to call SNPs in reads that mapped uniquely to one location on one haplotype, under stringent conditions, using a Q>30 filtering of both the Longranger mapping calls and FreeBayes variant calls. For Bcftools, filters allowed only homozygous ALT alleles (GT=“A”), as the reference assembly used was a concatenated diploid assembly instead of a haploid one. 1,862,877 variants were called. The resulting diploid assembly (Mar.3.4.6.p0p1_Q30Q30A.fasta) was split into the two haploid genomes and renamed (Mar.3.4.6.p0_Q30Q30A.fasta and Mar.3.4.6.p1_Q30Q30A.fasta). Commands for running of polishing and renaming of the assembly are available (LongrangerFreeBayesBcftoolsPolishing.txt).

Phase 1 was selected as the primary reference genome for annotation and further use, as it contained the first endogenous *Steamer* insertion site that was initially reported (*43*). This site is polymorphic in *Mya arenaria* populations and was not present in Phase 0.

#### RNA extraction and transcriptome assembly

Seven tissues from the reference animal (MELC-2E11) frozen at -80°C in RNAlater (Invitrogen, Waltham, MA) were used for RNA extraction (1, mantle; 2, foot; 3, siphon; 4, stomach; 5, adductor muscle; 6, gills; and 7, hemocytes). Solid tissues were homogenized with a disposable plastic mortar and pestle in liquid nitrogen before extraction with the Qiagen RNeasy kit (Qiagen, Hilden, Germany), eluting in 60 µL elution buffer. DNase I (2 µL, 2,000 U/ml, RNase-free, New England Biolabs, Ipswich, MA), 10× DNase buffer, and water was then added to a total of 100 µL, and the reaction was incubated for 1 h at room temperature. Then 250 µL ethanol was added and mixed by pipette, and it was added to a second Qiagen RNeasy column. The RNeasy protocol was followed, skipping the RW1 step, adding 500 µL RPE 2×, and eluting in 40 µL elution buffer. RNA samples (excluding the stomach due to possible contamination with RNA from clam food) were then sequenced (Genewiz, Leipzig, Germany).

RNAseq reads from the six tissues were concatenated to create single files for each read direction and used to assemble a transcriptome using Trinity (v2.8.5) (*68*):

➢ Trinity --seqType fq --max_memory 200G --CPU 16 --trimmomatic --full_cleanup --left MELC-2E11_R1_allfiles-cat.fastq.gz --right MELC-2E11_R2_allfiles-cat.fastq.gz

#### Genome annotation

Repeat elements in the genome assembly were called using RepeatModeler (v2.0) and masked repeat elements using RepeatMasker (v4.1.0) (*69*):

➢ RepeatModeler -database Mar.3.4.6.p1_Q30Q30A -pa 20 -LTRStruct
➢ RepeatMasker -pa 20 -lib $i-families.fa Mar.3.4.6.p1_Q30Q30A.fasta

The genome was annotated using MAKER (v2.31.10), exonerate (v2.2.0), and RepeatMasker (v4.1.0) (*69*), with two rounds of SNAP training using custom scripts, following previously established methods (*70*). The *Mya arenaria* transcriptome (as assembled above) was used as input into the MAKER annotation, along with the proteins identified from five well-annotated bivalve genomes (*Mytilus coruscus*, GCA_011752425.2; *Crassostrea virginica*, GCF_002022765.2; *Mizuhopecten yessoensis*, GCF_002113885.1; *Pecten maximus*, GCF_902652985.1; and *Crassostrea gigas*, GCF_902806645.1).

Putative gene identification was made by BLASTP search of the uniprot database (accessed 2021-03-02) and the proteins identified from the five well-annotated bivalve genomes concatenated into a single file (CgiCviMcoPmaMye_protein.fasta), using blast+ (v2.10.0). The top hit (with and e value <1e^-6^) was used for gene identification. Genes were labeled based on the most similar uniprot hit (if applicable with an e value <1e^-6^) and “-like” suffix or labeled as uncharacterized if only matching an uncharacterized bivalve gene or no gene at all. To account for multiple genes with the same uniprot hit, an additional numeric suffix was added to indicate additional hits to the same uniprot gene or uncharacterized genes (e.g. _1, _2, _3…).

➢ wget ftp://ftp.uniprot.org/pub/databases/uniprot/current_release/knowledgebase/complete/uniprot_sprot.fasta.gz
➢ gunzip uniprot_sprot.fasta.gz
➢ makeblastdb -in uniprot_sprot.fasta -out uniprot_sprot -dbtype prot
➢ blastp -query /home/metzgerm/MAKER_Mya/2020-09-11-Mar.3.4.6.p1-MAKER/snap02/2020-09-11_Mar_genome_snap02.all.maker.proteins.fasta -db uniprot_sprot -evalue 1e-6 -max_hsps 1 -max_target_seqs 1 -outfmt 6 -out 2020-09-11_Mar_genome_snap02.all.maker.proteins.fasta.blastp -num_threads 20
➢ makeblastdb -in CgiCviMcoPmaMye_protein.fasta -out CgiCviMcoPmaMye_protein -dbtype prot
➢ blastp -query /home/metzgerm/MAKER_Mya/2020-09-11-Mar.3.4.6.p1-MAKER/snap02/2020-09-11_Mar_genome_snap02.all.maker.proteins.fasta -db CgiCviMcoPmaMye_protein -evalue 1e-6 -max_hsps 1 -max_target_seqs 1 -outfmt 6 -out 2020-09-11_Mar_genome_snap02.all.maker.proteins.fasta.CgiCviMcoPmaMye_blastp - num_threads 20

#### Genome assembly summary statistics

Genome assembly statistics for the current and previous *Mya arenaria* assemblies can be found in **Supplementary Table 1**. Genome size, GC content, scaffold N50 and contig N50 were calculated using BBTools stats.sh (v38.86) (*71*). Repeat content was estimated by running RepeatMasker (v4.1.0) using the previously generated RepeatModeler repeat library. Benchmark of Universal Single Copy Orthologs (BUSCO) scores were calculated against the metazoa_odb10 database using BUSCO v3 (*22*).

### MarBTN genome sequence analysis

#### Sample collection and DNA extraction

MarBTN samples were collected from highly neoplastic clams from Maine and New York, USA and Prince Edward Island, Canada (**Fig. 1A and Supplementary Table 2)**. Hemolymph DNA from several samples of MarBTN have been previously reported (those collected 2009-2014) (*5, 43*), and remaining samples (those collected 2020-2021) were shipped live on ice from a seafood supplier in Maine. Hemolymph was drawn and screened for highly neoplastic animals (as described above), and genomic DNA was extracted using the same protocol previously used for other MarBTN samples (DNeasy Blood and Tissue Kit, Qiagen)(*5, 43*). Two healthy clams were collected and DNA extracted from siphon or mantle tissue as reported previously(*5*), in addition to the healthy reference individual described above. Previous reports of likely BTN in *M. arenaria* (**Figure 1A**, x’s) were collected from reports in which “disseminated neoplasia” or “hemic neoplasia” were diagnosed in *Mya arenaria* (*21, 32, 72–93*).

#### Whole genome sequencing and genome mapping

All samples were sequenced by Genewiz on an Illumina HiSeq. Healthy tissue and cancer hemolymph were sequenced using a full lane with a target read depth of 50× given the expected haploid genome size. Paired tissue samples for a subset of cancer samples were sequenced on a half lane with a target read depth of 30×. Illumina sequences were purged of optical duplicates using BBTools clumpify (v38.86)(*71*), trimmed using trimmomatic (v0.36) (*94*) with a read quality threshold of 20, and mapped to the reference genome using BWA-MEM (*95*) with default settings.

- 02_Illumina_data_processing/02_map_to_genome.sh
- 02_Illumina_data_processing/00_sampling_map.R
- 02_Illumina_data_processing/01a_dedupe_and_trim.sh
- 02_Illumina_data_processing/01b_dedupe_and_trim_newsamples.sh

#### SNV calling

SNVs and indels were called using somatypus (v1.3), a platypus-based variant calling pipeline designed for closely related cancer data without a paired normal sample, ideal for the analysis of transmissible cancer genomes (*11*). Variants were called as present in a healthy clam if they were called by somatypus and supported by more than 3 reads. For cancer samples, we used more stringent thresholds to eliminate contaminating host DNA from being called as cancer alleles. Paired host tissue samples proved to be too highly contaminated by cancer to be useful in eliminating host alleles and were only used as a downstream confirmation that we were eliminating host alleles with our read thresholds. Unlike the mammalian transmissible cancers, which form solid tumors and allow the collection of uncontaminated healthy host DNA, BTN disseminates into the tissues of the host as the cancer progresses, resulting in tissue samples that include significant BTN cells, often even more cancer cells than host cells, in late stages of the disease. However, we find DNA extracted from hemolymph to be so highly composed of BTN cells and so few host hemocytes (**Supplementary Fig. 2)**, that we were able to effectively remove host variants using these thresholds.

Sequencing resulted in a range of average sequencing depths of 57-90 at called SNV loci across samples. We normalized SNV calling thresholds to the average read depth for each individual sample, to avoid biasing calls in favor of more deeply sequenced samples. A variant was called as present in cancer if it was present in at least one cancer sample at a depth of greater than 1/6 the average read depth for that sample (9-15 reads). Given a variant passed that criteria for at least one cancer sample, it was called in any other cancer sample if it was present at a depth greater than 1/16 the average read depth for that sample (3-5 reads). These thresholds were chosen to minimize the calling of host alleles or mis-mapped reads as cancer alleles, while also preventing the exclusion of real cancer alleles from our variant set.

- 02_Illumina_data_processing/03_rename_for_somatypus.sh
- 02_Illumina_data_processing/04_run_somatypus.sh
- 03_SNV_analysis/02_initial_SNV_counts.R

### LOH region identification

To call genome regions where a haplotype was lost in one sub-lineage but retained by the other sub-lineage (termed LOH for loss of heterozygosity), we focused on SNVs for which we had high confidence that they came from the founder clam germline – those found in all cancer samples and at least one healthy clam. We calculated the allele frequencies for each of these SNVs in each cancer sample and flagged SNVs that were likely homozygous (above 0.8 frequency) for all samples in one sub-lineage, while heterozygous (less than 0.8) for all samples in the other sub-lineage. We included three representative samples from the USA sub-lineage in these calls (FFM-19G1, FFM-22F10 and NYTC-C9) so that calls were not biased by there being more USA samples than PEI. A region with SNVs transitioning to homozygous from heterozygous (the ancestral heterozygous state being captured in the other sub-lineage) would indicate regions that had lost a parental haplotype in the homozygous sub-lineage. We looked at sliding windows of 50 heterozygous SNVs across each scaffold and counted the number of SNVs that were heterozygous -> homozygous discordant for each sub-lineage. We found that windows with 10 or more discordant SNVs were the most effective for calling LOH regions (see below for validation of this threshold). We merged overlapping windows, for a total of 1,098 LOH windows for PEI (155 Mb, 12.8% of the genome) and 817 LOH windows for USA (98 Mb, 8.1% of the genome).

The amount of the genome that was called as in an LOH region was highly sensitive to the threshold of heterozygous -> homozygous discordant SNVs used to call LOH windows. We used two metrics to determine the best threshold, signature S mutation fraction and dN/dS. Both metrics are proxies for somatic mutations, with lower values corresponding to more founder variants and higher values corresponding to more somatic mutations. A higher threshold results in a high confidence in the LOH regions, but with missed true regions of LOH, resulting in more founder variants in regions called as non-LOH, while a lower threshold results in over-calling of LOH regions but with less founder variants in the regions called as non-LOH. We tested the calling of LOH regions as described above for all possible thresholds between 0 and 50 SNVs in the 50 heterozygous SNV window. We then divided high confidence somatic mutations into the regions called in these test sets as LOH and non-LOH. We then calculated signature S mutation fraction and dN/dS ratio for each and plotted the values against the threshold used for the test calling (**Supplementary Fig. 3**). Over-calling of LOH drops dramatically before flattening out around 10/50 SNVs, while missed LOH appears to rise consistently as the threshold is increased. Overall, a threshold of 10/50 SNVs maximizes the difference between somatic mutations in non-LOH vs LOH regions and was used in all other analyses to call LOH regions, so that they could be excluded from somatic mutation analysis.

- 03_SNV_analysis/04a_LOH_calling_upstream.R
- 03_SNV_analysis/04b_LOH_merge_and_helmsman.sh
- 03_SNV_analysis/04c_exclude_LOH_SNVs.sh
- 03_SNV_analysis/06b_run_dNdS.sh
- 03_SNV_analysis/07_LOH_threshold_validation.R

#### MarBTN phylogeny

The phylogeny in **Fig. 1E** is a neighbor-joining tree was built using R package “ape” (v5.5) (*96*) from a table of pairwise SNV differences between cancer samples. SNVs found in any healthy clams were excluded prior to this analysis, since nearly all those SNVs were likely present in the founder clam. SNVs in LOH regions were also excluded. The reference genome was added to serve as the tree root (by definition, the reference genome does not have any of the called SNVs). Bootstrapping revealed high confidence (100/100) for the PEI and USA sub-lineage nodes, but low confidence for all other nodes (<20/100).

- 03_SNV_analysis/01_pairwise_phylogeny.sh
- 03_SNV_analysis/02_initial_SNV_counts.R

#### Mutational signature extraction and fitting

We categorized SNVs into 25 bins based on which samples they were found in (see **Supplementary Fig. 20** or code reference below). We further divided each SNV bin by annotated genome regions (full genome, genes, exons, CDS, 5’UTR, 3’UTR). We used Helmsman (v1.5.2) (*97*) to count SNVs for each bin in their trinucleotide context, and R package “Biostrings” (v2.54.0) to count trinucleotide opportunities in each genome region. We performed *de novo* signature extraction on this data set using R package “sigfit” (v2.0.0) (*98*), correcting for opportunities in each genome region. The unbiased estimate for the best number of signatures to fit our data was 3, though extracting 4 signatures revealed a signature of unmistakable resemblance to COSMIC signature 1 (CpG>TpG) so we proceeded with 4 signatures. SNV bins were then reanalyzed with these 4 signatures, again correcting for mutational opportunities, to reveal the fraction of SNVs in each category that could be attributed to each signature.

- 03_SNV_analysis/03a_sig_extraction_upstream.R
- 03_SNV_analysis/03b_sig_extraction_count_trinuc.sh
- 03_SNV_analysis/03c_sig_extraction_fitting_sigfit.R
- 01_Genome_assembly/03a_trinucleotide_counting.sh
- 01_Genome_assembly/03b_trinucleotide_counting.R

#### Cancer dating and estimation of total somatic SNVs

To estimate the age of the MarBTN lineage we only wanted to consider likely somatic mutations, so we excluded regions that were called as LOH in either sub-lineage from these analyses (as founder SNVs in a region lost in one sub-lineage would appear to be unique to the other sub-lineage and could be falsely considered to be late somatic SNVs if those regions were not removed). We then filtered remaining SNVs from each MarBTN sample to remove any SNVs that were found in a healthy clam or the other sub-lineage using the same thresholds as described above (SNV calling). The remaining SNVs for each sample should be somatic mutations which occurred since the time the two sub-lineages diverged (since the most recent common ancestor, or MRCA). We counted the number of mutations in trinucleotide contexts using Helmsman (*97*) for each MarBTN sample and fit this to our *de novo* extracted mutational signatures to estimate contributions of each of the 4 signatures. We then performed a linear regression of the mutation count attributed to each signature for each sample against the date the sample was collected (**Fig 2C** and **Supplementary Fig. 8)**. Signature S was the signature which best fit with time, perhaps due in part to it being somatic mutation-specific and therefore excluding false positive SNVs that may have made it through the filters due to missed LOH or other errors.

The x-intercept of the regression calculated above indicates the age of the most recent common ancestor of the two sub-lineages (i.e., when mutation count separating them equals zero). To estimate the total age of the cancer, we first estimated the number of somatic signature S mutations in the trunk of the MarBTN lineage: SNVs shared by all MarBTN samples. We continued to exclude LOH regions, to work with the same region for which the mutation rate was calculated, and to exclude SNVs shared with any healthy clams, which are presumably founder variants. Somatic mutations in non-LOH regions with copy number >1 would have been heterozygous when they occurred, so we filtered for SNVs with an average allele frequency under 0.8 across the 8 MarBTN samples. For comparison, we also analyzed the following SNV bins: likely homozygous SNVs (those with an average allele frequency over 0.8); SNVs in healthy clams (from each of the three healthy clams individually as well as all SNVs found in any healthy clam); SNVs found in all cancer samples and shared with a healthy sample; and SNVs found in all samples in one sub-lineage but not the other sub-lineage or healthy clams (high confidence somatic mutations). We counted mutations in their trinucleotide contexts and fit the 4 *de novo* extracted signatures as described previously. The fraction of SNVs found in healthy clams attributable to signature S was taken to be the baseline signature S fraction (0.025). The signature S mutation fraction was near this baseline for individual healthy clam SNVs and for likely founder variants – those found in all MarBTN samples and either shared with a healthy clam or not shared with a healthy clam but homozygous. Heterozygous SNVs found in all MarBTN samples but no healthy samples had noticeably higher signature S fraction (0.056). The difference between this fraction and the baseline was taken to be from signature S mutations in the early somatic evolution of the MarBTN lineage (0.056-0.025=0.031). This fraction is equivalent to 53,350 mutations, or 126 years (95% CI: 89-224) by the previous signature S mutation rate calculation. This confidence interval was determined solely from uncertainty in the mutation rate since error estimates from signature fitting with sigfit were negligible in comparison.

Using the signature S mutation estimations above, we then estimated the total number of somatic mutations in each of the MarBTN sub-lineages, which would be a combination of mutations occurring post-MRCA (high confidence somatic mutations) and mutations in the lineage trunk. To first estimate somatic mutations in the trunk, we assumed that somatic signature S mutation fraction had remained steady since oncogenesis at 0.48 (based on high confidence somatic mutations in the two sub-lineages). Given the following equations describing the total number of mutations being comprised of a fraction of founder SNVs and a fraction of somatic SNVs (a) and that each fraction has a known percentage of mutations due to SigS (b), we then solve for Fraction_somatic_ (c-e):

a. Fraction_founder_ = 1 – Fraction_somatic_
b. SigS_observed_(heterozygous all BTN, no healthy)=Fraction_somatic_*SigS_somatic_+Fraction_founder_*SigS_founder_
c. SigS_observed_=Fraction_somatic_*SigS_somatic_+ (1 - Fraction_somatic_)*SigS_founder_
d. Fraction_somatic_ = (SigS_observed_ - SigS_founder_) / (SigS_somatic_ - SigS_founder_)
e. Fraction_somatic_ = (0.056-0.025) / (0.48-0.025) = 0.068

… which is equivalent to 116,765 somatic mutations. We then added high confidence somatic SNVs unique to each sub-lineage (those present in all samples from that sub-lineage but none in the other sub-lineage or healthy clams: 431,987 for PEI and 443,424 for USA) and corrected for genome size of the non-LOH portion of the clam genome to get mutation density estimates for each sub-lineage. Note that although we can estimate total mutation count in the lineage trunk, we cannot differentiate individual SNVs as somatic mutations or founder variants.

- 03_SNV_analysis/05a_cancer_dating_prelim.R
- 03_SNV_analysis/05b_cancer_dating_helmsman.sh
- 03_SNV_analysis/05c_cancer_dating_regression.R
- 03_SNV_analysis/05d_cancer_dating_trunk.R

#### dN/dS

We calculated global dN/dS, the overall ratio across all genes in the genome, after filtering out the following SNV subsets:

- SNVs found in all healthy clams
- SNVs found in any healthy clam
- SNVs unique to each of the three healthy clams
- SNVs in all MarBTN samples and shared with a healthy clam
- SNVs in all MarBTN samples and not found in any healthy clams in our data set
- SNVs found in all samples for each sub-lineage, but not found in the other sub-lineage or healthy clams. This resulted in three subsets: USA, PEI and SNVs from each sub-lineage combined. These were further filtered to include only SNVs outside called LOH regions.

We ran R package “dNdScv” (v0.0.1.0)(*35*) for call dN/dS for each SNV subset. dNdScv is designed to quantify selection during somatic evolution and corrects for trinucleotide context-dependent biases to estimate a dN/dS ratio normalized to the expected ratio for each gene or the entire genome. dNdScv is designed to be run on datasets of many samples, but in our case we ran it individually on the above SNV subsets. We ran dNdScv with default settings except for setting max_coding_muts_per_sample and max_muts_per_gene_per_sample to 1 billion each, effectively removing these maximum settings, which were designed for conventional cancers. We calculated global dN/dS across the whole genome, including all annotated genes. Likely somatic SNVs show largely neutral global dN/dS (0.98, 95%CI: 0.94-1.02), indicating that there is minimal contamination from founder variants, which are assumed to have been predominantly under negative selection, as seen in healthy clams SNVs.

We also calculated dN/dS for individual genes to search for signals of positive or negative selection. We filtered for genes under significantly positive or negative selection (corrected p-value < 0.05). For the five hits generated when dN/dS was run for somatic mutations, we performed an NCBI blastp query for each of these genes, getting hits for three of the five genes. We checked each gene visually/manually using IGV, noting that in each case nearly all SNVs appear to be on a single haplotype. We calculated the dN/dS for SNVs found in any healthy clam for each of these 5 genes, and none were under significantly positive selection in the observed healthy clam genomes (this would have been expected if these hits were due to missed founder variants in a gene under positive selection in the healthy clam population). Results and notes for each gene are summarized in **Supplementary Table 3**.

- 03_SNV_analysis/06a_bin_for_dNdS.R
- 03_SNV_analysis/06b_run_dNdS.sh
- 03_SNV_analysis/06c_dNdS_outputs.R

#### Copy number calling

Most cancer copy number calling tools rely on having paired tissue samples, so we developed a custom copy number calling script based on read depth relative to the clam used to build the reference genome (MELC-2E11), with the assumption that this reference clam is diploid. First we used R package “cn.mops” (v1.32.0) (*99*) to divide the genome into 1 kB windows and count the number of reads mapping to each window for each of the samples: healthy (3) and MarBTN (8). Any window with low mapping in the reference clam (less than ¼ the average read depth) was excluded from calling as a low-mapping region. Read depth for each window for each non-reference sample was then divided by the reference clam read depth for that window to normalize. Each window was then divided by the average read depth for that sample to yield a log2 read depth score. We then calculated the median log2 read depth for every 100 1 kB windows to form larger windows of 100 kB.

We then wanted to convert log2 read depth to copy number without prior knowledge of the average ploidy for each sample. We observed distinct peaks corresponding to copy number integers when we plotted histograms of log2 read depth scores genome-wide for each sample. We chose the best fitting average ploidy value for each sample (the value which lined up copy number calls with integer values when multiplied by 2^log2_score^). This was 3.6 for PEI MarBTN samples, 3.3 for USA MarBTN samples, and 1.9 for non-reference healthy samples. Note that since healthy samples should be diploid, an average ploidy just under 2 is expected, given read mapping will be slightly less efficient for non-reference clams relative to the reference clam (whose reads are mapped to a reference genome built from itself). We multiplied log2 read depth scores by this value to get copy number estimate for each 100 kB window for each sample. Observing close agreement between the samples within each sub-lineage (**Supplementary Fig. 10**), we calculated the average copy number calls for each sub-lineage. Finally, we smoothed copy number calls in 1 Mb windows to minimize noise in final calls. For each 100 kB window we calculated the standard deviation for the preceding ten 100 kB windows, the following ten 100 kB windows and the surrounding ten 100 kB windows (five 100 kB windows on either side). We replaced the copy number call with the median of the 1 Mb window with the smallest standard deviation, provided the standard deviation was small, defined as less than 1 on the ploidy scale. If the standard deviation was larger than 1 for all windows, we left the original unsmoothed copy number. Finally, we rounded all calls to the closest integer value for the final copy number call for each 100 kB window. However, we kept the unrounded calls for the purpose of visualizing error in our figures (**Fig. 3B** and grey bars in **Fig. 3A**).

To validate our copy number calls, which were based solely on read depth, we used variant allele frequency of somatic mutations. If calls are correct, genome regions that are of a particular copy number should exclusively have certain allele frequencies, such as 0.33/0.67 for CN3 regions or 0.25/0.5/0.75 for CN4 regions. We calculated variant allele frequencies for high confidence somatic mutations, some of which likely occurred after copy number alteration events and therefore should have a frequency distribution peak around the low frequency value (e.g 0.33 for CN3 or 0.25 for CN4). We separated SNVs specific to each sub-lineage based on the copy number calls at their locations using bedtools (v2.29.1) (*100*) and calculated average variant allele frequency across each ploidy level in all of the samples. A plot of the variant allele frequency distribution shows that the major peak corresponds to the expected frequency for each copy number bin. There is evidence of some off-target peaks indicating some degree of error in these copy number calls (for example, 0.5 peak in CN3, 0.33 peak in CN2, 0.5 or less in CN1). Some of these peaks indicate regions that are called as lower copy number than the true value (e.g. 0.33 peak in CN2, 0.5 or less in CN1), which is likely due to sequence polymorphism leading to lower mapping than expected. Other off-target peaks, particularly those indicating copy number is called too high, may be due to other causes, such as the confounding effects of repetitive elements in the genome. Overall, copy-number-specific the variant allele frequencies support the conclusion that this copy number calling strategy is accurate and that much of the MarBTN genome has increased in copy number from its diploid founder ancestor.

- 04_CNV_and_SV_analysis/01_CNV_calling.R
- 04_CNV_and_SV_analysis/02_SNVs_by_CNV.sh
- 04_CNV_and_SV_analysis/03_SNV_freq_by_CNV.R

#### Structural variant and telomere calling

We used Delly (v0.8.5) (*101*) to call deletions, small (<100 bp) insertions, tandem duplications, inversions, and translocations in each sample individually from split read mapping. Delly is sensitive to read depth, so we subsampled all sample sequences to only include 600,000,000 reads (which is a lower count than the lowest sequenced sample) prior to running Delly using “samtools view -s”. We only considered SVs supported by reads mapping to precise breakpoints in the genome. We used default settings, except for setting a minimum paired end read mapping quality threshold to 30 to minimize false positives. We merged all called SVs into a single file based on shared breakpoints. We removed SVs called in the reference clam from all samples and compared the number of each SV type and size of each intra-chromosomal SV type. To narrow in on high confidence somatic SVs we then filtered out SVs found in any healthy clam or the opposite sub-lineage from each sample (similar to our approach for identifying somatic SNVs) and compared number and size of SVs. To compare SV counts between healthy/MarBTN and USA/PEI, we used a two-sample t-test (unequal variance) and to compare sizes we used a two-sample Wilcoxon signed-rank test.

We used telseq (v0.0.2) (*102*) using default settings to estimate telomere lengths.

- 04_CNV_and_SV_analysis/04_SV_calling_delly.sh
- 04_CNV_and_SV_analysis/05_SV_analysis.R
- 04_CNV_and_SV_analysis/06a_telomeres.sh
- 04_CNV_and_SV_analysis/06b_telomeres.R

#### Identifying *Steamer* insertion sites

We called *Steamer* insertion sites in all samples via a custom pipeline which uses split reads that map to both the reference genome and *Steamer* itself. First, we used BWA-MEM (*95*) to map reads for each sample to the 177 bp *Steamer* long terminal repeat (LTR), a sequence that flanks either side of the internal coding sequence of LTR-retrotransposons (*43*). For all reads that mapped, we extracted just the externally flanking portion of each read, discarding reads that extended into the internal sequence of *Steamer* and discarding the portion of each read that mapped to the *Steamer* LTR. We then mapped these flanking portions to the reference genome, keeping only reads that mapped to a single location in the genome with high confidence (MAPQ score ≥ 30). For reads mapping to the genome with lower confidence (MAPQ score < 30), we rematched each flanking read with its pair and re-mapped to the genome using bwa sampe(*59*), and if the mapping of the flanking fragment together with its mate generated a MAPQ score ≥ 30 then it was included as a specific mapped read. Finally, we took all flanking reads that did not map to the genome with high confidence in either step and mapped them to the RepeatModeler2-generated repeat library, since many flanking reads that do not map to a specific site in the genome are likely to be in repetitive regions. We then generated a BED file format for each flanking read mapped to the genome or repeat library, calling each *Steamer* site by its 5 bp target site duplication, which is generated upon insertion and means that upstream and downstream flanking reads will overlap by 5 bp, and whether it was forward or reverse-face relative to the mapped chromosome.

We merged reads by their mapped locations to get the total number of upstream and downstream flanking reads supporting each insertion site, keeping all sites supported by at least five total flanking reads or at least one each of upstream and downstream flanking reads. We corrected for the six *Steamer* insertions that exist in the reference genome, which would otherwise result in upstream and downstream reads mapping 4.7 kB apart if that insertion is present in a sample (before and after the *Steamer* copy in the reference genome), so that upstream and downstream reads were still counted as in support of the same insertion. We then merged these insertion calls for each sample into a single table, which we used to count shared insertion sites between samples (e.g. all MarBTN, PEI only, USA only) and build a phylogeny using pairwise differences using R package “ape” (*96*). We estimated total read depth at *Steamer* insertion sites by averaging the read depth 10 bp before and10 bp after the 5 bp target site duplication, only considering reads with MAPQ ≥ 30. We then estimated insertion allele frequency at each site by dividing the number of *Steamer* insertion supporting reads by the total read depth.

We noticed a bias for ATG in positions 7-9 in both our upstream and downstream *Steamer* flanking reads. To investigate this bias, we extracted the 35 bp surrounding each *Steamer* insertion sites from the reference genome (15 bp upstream, 5 bp target site duplication, and 15 bp downstream) using bedtools getfasta(*100*). We then counted the number of occurrences of each nucleotide at each position, normalized by the GC content of the genome (35%), and created logo plots using ggseqlogo (*103*). This bias held whether we looked at *Steamer* sites across all samples, just cancer samples, sites shared by all cancer samples, sites unique to the USA sub-lineage, and sites unique to the PEI sub-lineage. For sites found in any cancer sample, we also counted the number of sites that had an ATG in positions 7-9 upstream, downstream (note ATG in read in reverse is CAT), and both upstream and downstream. Compared to the frequency expected based on the frequency of ATG in the genome (2.2% of trinucleotides), these sites were 8.5, 7.4, and 44.6 times more frequent than expected by chance, respectively.

To investigate where *Steamer* inserted relative to genes, we found the closest gene to each insertion site using bedtools closest (*100*), excluding insertion sites within genes. There was a noticeable bias in the 1-2 kB upstream genes (**Supplementary Fig. 21A**). To ensure this was not due to read mapping bias, we generated a similar plot based on whole genome sequence read mapping by mapping 0.1% of MELC-2E11 reads to the genome and treating the first 5 bp as a *Steamer* insertion site. This test set did not display this bias (**Supplementary Fig. 21B**). We then counted the number of *Steamer* insertions in annotated regions in the genome (genes, coding sequences, 5’UTR and 3’UTR) in addition to the 1 kB regions upstream of annotated gene regions. We then normalized for both the size of those portions of the genome and how likely reads were to map to these regions (to correct for biases that might skew insertions toward more mappable portions of the genome), yielding the plot found in Figure 4C.

To see whether these genes might be more likely to be cancer-associated, we conducted a blastp search of predicted intact *M. arenaria* gene models for the 729 cancer-associated genes from the COSMIC database, which generated hits of e-value>1e-6 in 14% of *Mya* genes. We then compared the number of *Steamer* insertions that intersect with these genes. In the absence of selection for insertion near these genes we would expect 14% of *Steamer* insertions to intersect with these genes. Observed versus expected insertions were compared with a Chi-squared test. See **Supplementary Fig. 14** for results.

- 05_TE_analysis/01_identify_steamer_in_ref_genome.sh
- 05_TE_analysis/02_steamer_calling_pipeline.sh
- 05_TE_analysis/03_steamer_downstream_analysis.R
- 05_TE_analysis/04a_steamer_ATG_bias.sh
- 05_TE_analysis/04b_steamer_ATG_bias.R
- 05_TE_analysis/05a_steamer_upstream_bias.sh
- 05_TE_analysis/05b_steamer_upstream_bias.R
- 05_TE_analysis/05c_steamer_upstream_cosmic_bias.sh

#### TE copy number analysis

We did not observe *Steamer* in our RepeatModeler run on the reference genome, likely due to it being present at low copy number and thus not clearing the threshold to be called as a repeat element. In order to capture other repeat elements like *Steamer* that might be high copy number in MarBTN but low in the reference genome, we also ran REPdenovo (*104*), a repeat element identifier that can be run on raw WGS data, as opposed to the assembled genome required for RepeatModeler. We ran REPdenovo on the healthy reference clam (MELC-2E11), a USA MarBTN sample (MELC-A11) and a PEI MarBTN sample (PEI-DN08) to capture repeat elements at high copy number in either sub-lineage, as well as a healthy clam to control for biasing repeat element identification towards MarBTN. We then ran RepeatClassifier, a component of RepeatModeler used for classifying repeats based on sequences, on the output repeat elements.

To generate a consensus repeat library, we used CD-HIT (v4.8.1) (*105*) to merge the libraries generated from the RepeatModeler and REPdenovo runs, using the same CD-HIT settings as those used by RepeatModeler itself to merge repeats with greater than 80% identity (-aS 0.8 -c 0.8 -g 1 -G 0 -A 80 -M 10000). We then used BWA-MEM to map reads from each sample to the repeat library and calculated the average read depth across each repeat element. We then normalized by read depth across the genome, calculated previously, to control for variation in sequencing depth between sequencing runs, to yield an estimate of the number of copies of each repeat element in each sample. Note that this copy number is relative to the haploid genome for all samples, so ploidy differences between our samples should not affect our copy number comparisons.

For each repeat element, we calculated the average copy number among our three healthy clams, eight MarBTN samples, and each MarBTN sub-lineage individually (five USA samples and three PEI samples). We calculated the ratio of copies in healthy clams versus MarBTN samples and PEI sub-lineage versus the USA sub-lineage, followed by a two-sample t-test to calculate the significance of each difference. We removed repeats with less than 1 copy in any sample, as these likely represent TEs that are only present in a subset of the clam population and would yield a highly significant difference simply due to the absence in some samples and presence in others. The remaining elements are plotted in the volcano plot in **Fig.4D** and **Supplementary Fig. 15**. We calculated a significance threshold using the Bonferroni correction for multiple tests for a corrected p<0.05. We additionally divided and plotted the data set by repeat type classified by RepeatClassifier (DNA transposon, LTR, LINE, rolling circle, rRNA, simple repeat, SINE, snRNA, or tRNA). We performed Chi-squared tests to determine whether certain elements were higher copy number in one group versus another. We also note that although we can conclude differences in copy number, many differences may be due to variation between the founder clam and the three healthy clams sequenced in this study, as opposed to being due to somatic expansions. The magnitude of repeat expansions may be overestimated, since we are comparing an average from three difference clams to an average from eight samples of a clonal lineage. However, the strong skew towards more copies in MarBTN compared to healthy clams indicates that either A) the founder clam had more copies of many TEs than the healthy animals sequenced here or B) many TEs have increased their copy number through somatic expansion.

- 05_TE_analysis/06a_REPdenovo.sh
- 05_TE_analysis/06b_merge_repeats_and_maps_reads.sh
- 05_TE_analysis/07_TE_coverage_analysis.R

#### Mitochondrial analysis

We mapped each whole genome sequenced sample to the previously published mitochondrial genome (*42*) using BWA-MEM (*95*). We then ran somatypus (*11*) using default settings to call SNVs and indels. We excluded SNVs around the multi-copy region in positions 12,060-12,971. We did not see evidence of heteroplasmy outside this region, so an SNV was counted as present if it was present in a sample at >0.5 VAF. To infer relatedness of mitochondrial genotypes we built a neighbor-joining tree, as done for genome SNVs and *Steamer* insertions, from a table of pairwise SNV differences between all samples, including the reference mitochondrial genome as the root of the tree.

We used mean allele frequency of mitochondrial SNVs as a proxy for cancer isolate purity and host tissue purity, as shown in **Supplementary Fig. 2**. For healthy clams, mean allele frequencies were slightly below 100% likely due to sequencing, mapping, or contamination errors, and yield a maximal value for “pure” target DNA. For cancer samples, mean allele frequencies were slightly lower, attributed to the presence of host clam DNA, but remained >97%. For samples for which paired tissue sequencing existed (5 of 8 cancer samples), we used the mean allele frequency of cancer-specific mitochondrial SNVs to estimate contamination of tissue by cancer DNA. Some samples contain a high amount of cancer DNA, making genome-wide differentiation between host and cancer SNVs difficult in tissue and leading us not to include paired tissue DNA in our analyses.

To look at mutational biases, we included 12 possible single-nucleotide substitution types rather than the traditional 6, since the heavy/light strand differences of mtDNA result in unequal C/G and A/T in the forward or reverse direction (in fwd direction: A=0.29%, T=0.37%, C=0.12%, G=0.23%). We counted SNVs of each substitution type for SNVs found in healthy clams (39), shared among all MarBTN samples but not found in healthy clams (13), those found in all samples of the USA (21) or PEI (26) sub-lineages, and all high confidence somatic mutations (50: those found in only a subset of MarBTN samples). We also calculated the expected number of substitutions of each type based on the nucleotide content of the mitochondrial genome assuming no mutational biases for comparison.

We used dndscv (*35*) as described previously to calculate global dN/dS in the mitochondrial genome. We calculated dN/dS for SNVs found in healthy clams, SNVs shared among all cancer samples but not found in healthy clams, and high confidence somatic mutations (i.e. those found in just the USA or PEI sub-lineages). 95% confidence intervals from dndscv are quite large due to the small number of coding mitochondrial mutations in our samples used for this calculation.

We calculated read depth at each position using samtools depth (*61*). To estimate the number of copies of the D-loop region, we calculated the average read depth in positions 12,300-12,500 relative the average read depth across the full mitochondrial genome excluding that region. This region was chosen because it is within the multi-copy D-loop region but should not have reads that border the duplication breakpoint or the insertion that is only present in some copies and may cause errors in amplification due to its G-rich sequence. Copy numbers were compared between the groups using a two-sample t-test (unequal variance).

The new reference mitochondrial genome was assembled by taking the previously published mitogenome reference with only a single, collapsed copy of the repeated D-loop region (NC_024738.1), replacing the 696 bp putative repetitive region (12,163-12,857) with a gap of 3×696 Ns, and running PBJelly to fill the gap using PacBio long reads. PBJelly was run to gap-fill the scaffolded assembly using pbsuite (*66*) (https://github.com/esrice/PBJelly) using using blasr v5.1, networkx 2.2, and Python 2.7 as above, with the protocol file Protocol_MELC.xml. Only captured gaps were filled (no inter-scaffold gaps) using the option “--capturedOnly” during the “support” step. PBJelly was run with the commands:

➢ Jelly.py setup Protocol_ MELCmtmultifastq.xml
➢ Jelly.py mapping Protocol_ MELCmtmultifastq.xml
➢ Jelly.py support Protocol_ MELCmtmultifastq.xml -x “--capturedOnly”
➢ Jelly.py extraction Protocol_ MELCmtmultifastq.xml
➢ Jelly.py assembly Protocol_ MELCmtmultifastq.xml -x “--nproc=20”
➢ Jelly.py output Protocol_ MELCmtmultifastq.xml

Polishing of the mitochondrial genome assembly was done with Arrow (using pbsuite as described above and pbbioconda-0.0.5 with python 3.7). First, the PBJelly output was renamed to PBJelly_mt_genome.fasta, and polishing was run using the commands:

➢ module load pbsuite/esrice
➢ pbalign --verbose --nproc 40 /home/metzgerm/MELC-2E11/Marenaria.3.2_bam.fofn PBJelly_mt_genome.fasta MELC-mtalignedall.bam 2>&1 | tee pbalign_stderrout.txt
➢ module load conda/4.7.10_py3.d/genomicconsensus/2.3.3
➢ samtools faidx PBJelly_mt_genome.fasta
➢ arrow --verbose --annotateGFF --reportEffectiveCoverage -j 40 MELC-mtalignedall.bam -r PBJelly_mt_genome.fasta -o MELC-2E11mtvariants.gff -o MELC-2E11mtconsensus.fasta -o MELC-2E11mtconsensus.fastq 2>&1 | tee arrow_stderrout.txt

The polished mitogenome alignment with the completely assembled repeat region (MELC-2E11mtconsensus.fasta) was renamed to mtGenome_PBJelly_polished.fasta. We confirmed the presence of a D-loop tandem duplication in a healthy clam using inverse PCR (**Supplementary Fig. 22)**, with outward facing primers that would only amplify if the copies or the region are in tandem (**Supplementary Table 4**). Amplifcation of the products of these inverse primers confirms tandem duplication of the region. However, amplicon sizes from primers spanning the D-loop support a single copy of the D-loop. Additionally, the inverse primers spanning the G-rich insertion has a dim band at expected size, but two brighter bands at smaller sizes. Given the highly G-rich region, it is likely that when primers spanning the D-loop are used that the PCR products are recombining to lose the extra copies, with selection in the PCR reaction favoring removal of the G-rich stretch that interferes with amplification. Given all samples in this study support the presence of tandem D-loop repeats, it is possible that the clam used for the previously published mitochondrial genome that contains a single D-loop copy (*42*) may have also been multi-copy and missed due to short-read sequences and recombination during cloning to resolve gaps in the mitochondrial genome.

- 06_Mito_analysis/00_create_coding_dndscv_input.sh
- 06_Mito_analysis/01_mapping_and_SNV_calling.sh
- 06_Mito_analysis/02_host contamination.R
- 06_Mito_analysis/03_dloop_coverage.R
- 06_Mito_analysis/04_SNV_analysis.R

