## Supplementary Figures and Tables for "Centuries of genome instability and evolution in soft-shell clam transmissible cancer"

### SUPPLEMENTAL FIGURES

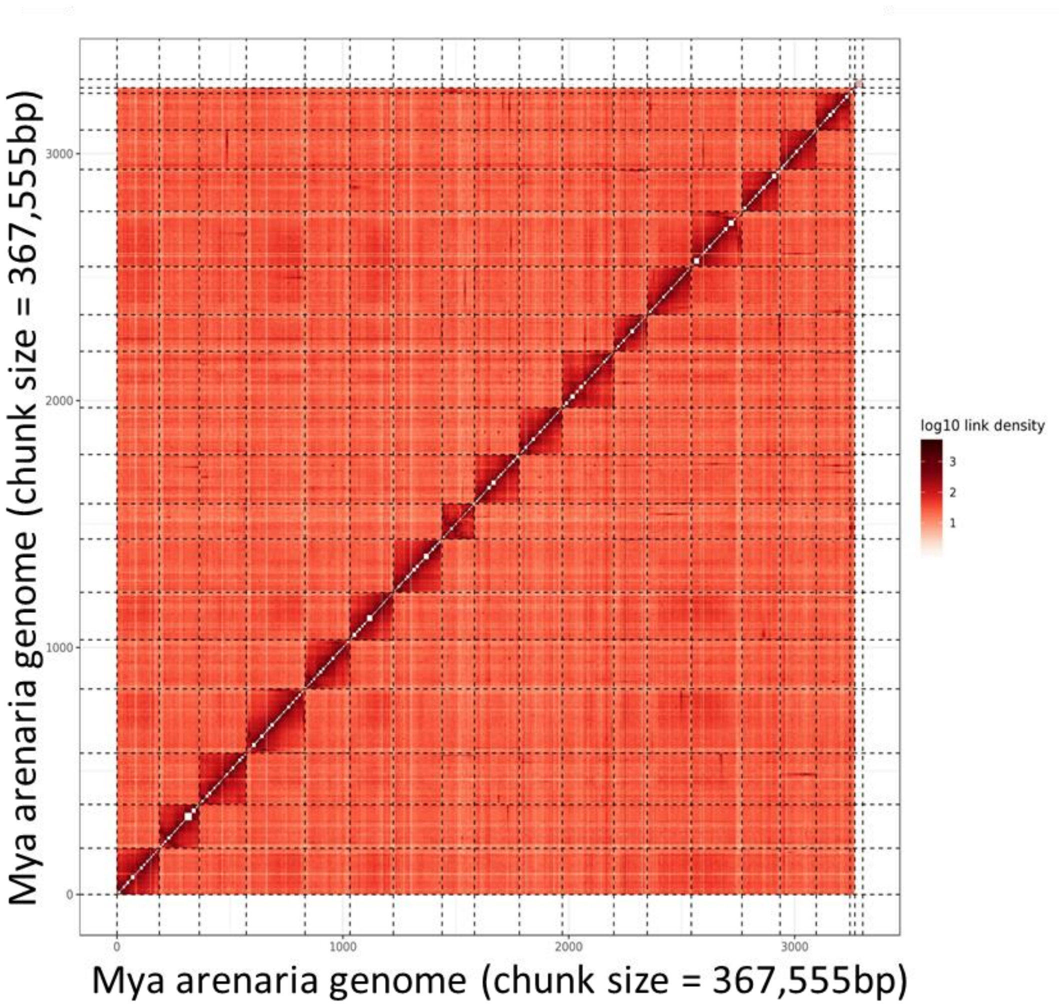


**Supplementary Figure 1: Hi-C scaffolding yields 17 presumptive chromosomes**
Heatmap of link density from Hi-C scaffolding, showing proximity of DNA segments in physical space across sequenced cells and clustering by chromosome. Results clearly yielded 17 scaffolds, matching the expected number of chromosomes in *M. arenaria*.

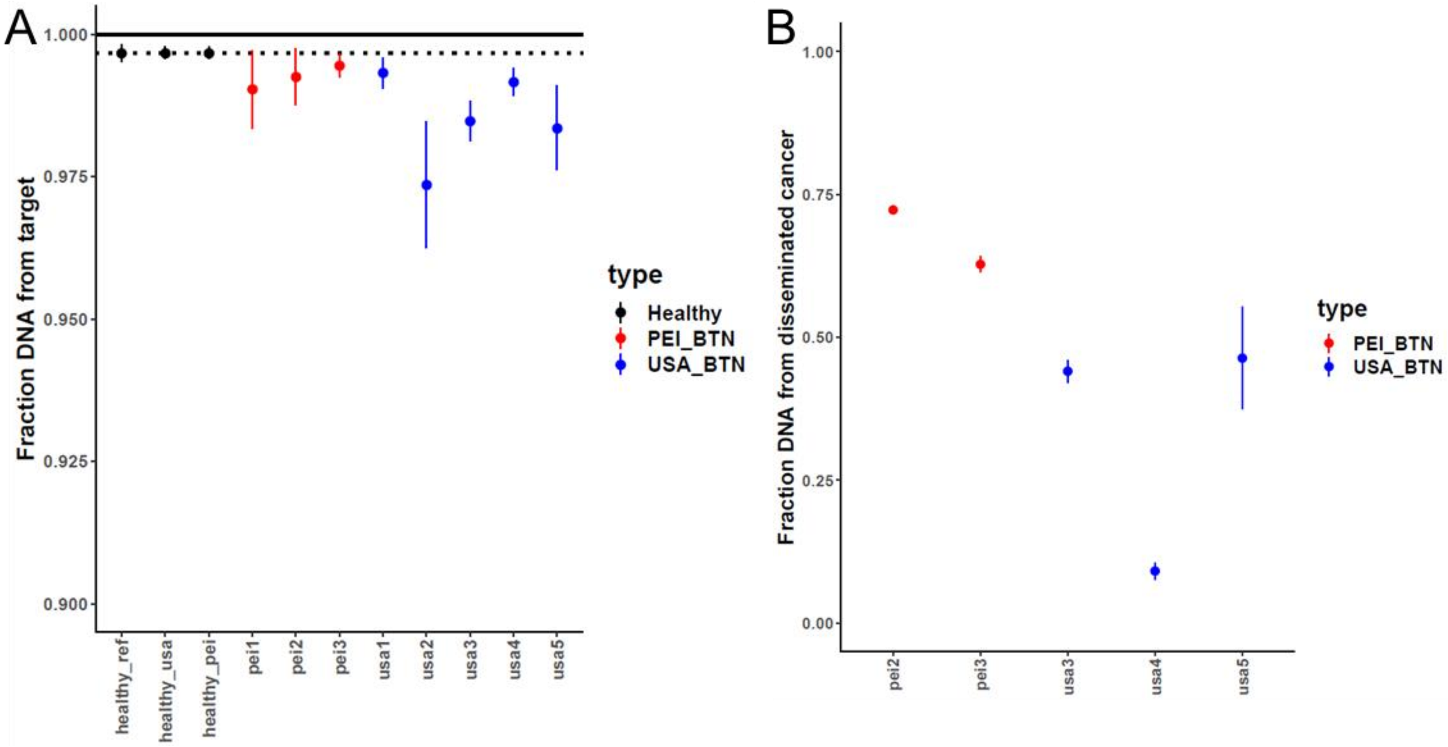


**Supplementary Figure 2: Minimal host DNA is found in cancer hemolymph samples, while high cancer DNA is found in some tissues**

**(A)** We estimated the percent of sequenced DNA attributed to targeted sample from the mean allele frequency across mitochondrial SNVs for each sample (number of loci: 13-21 for healthy clams, 46-53 for MarBTN samples). For healthy hemocyte samples (first three, black points), these values are slightly below 100% likely due to sequencing, mapping, or contamination errors, and yield a maximal value for "pure" target DNA. For cancer samples (last 8, red and blue points), the remaining drop in target DNA is attributed to the presence of host clam DNA and remains <3% in all eight analyzed samples. **(B)** We also extracted DNA from tissue samples and estimated the fraction of cancer DNA disseminated into tissue using the allele frequency of cancer-specific mitochondrial SNVs. We only sequenced tissue samples from five of eight MarBTN-infected clams in this study. Tissue samples contain variable, and in some cases quite high, fractions of cancer DNA. This made genome-wide differentiation between host and cancer SNVs difficult in tissue and lead us to not include paired tissue DNA in our analyses, instead relying on variant calling thresholds to eliminate host variants from our cancer variant calling pipelines. Error bars represent 95% confidence intervals.

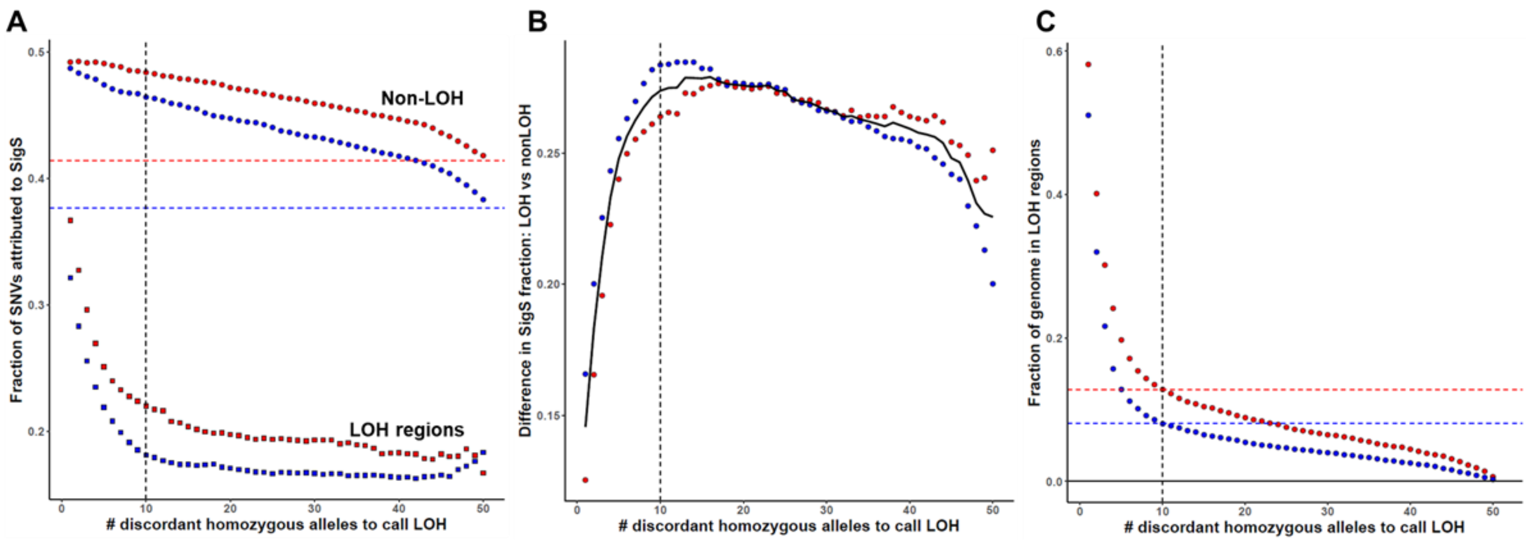


**Supplementary Figure 3: Calibrating LOH calling thresholds**

**(A)** We used various thresholds of stringency to call LOH across the genomes of each sub-lineage based on the number of shared SNVs that were homozygous in one sub-lineage but heterozygous in the other across a window of 50 SNVs (x-axis). After calling LOH, we calculated the fraction of likely somatic mutations attributed to signature S in LOH (squares) and non-LOH (circles) (y-axis). Values are shown separately for the BTN subgroups from USA (blue) and PEI (red). Vertical dashed line indicates the threshold used for LOH-calling. Horizontal dashed lines indicated baseline signature S fractions without LOH region removal. **(B)** Plot of the difference between non-LOH and LOH regions as shown in (A) (calculated by subtracting the square from the circle). Black line shows the average difference, which peaks around the threshold used (10). **(C)** Proportion of the genome that is called LOH for each sub-lineage based on calling threshold. Dashed lines indicate the fraction of the genome called as LOH for each sub-lineage for the final threshold used.

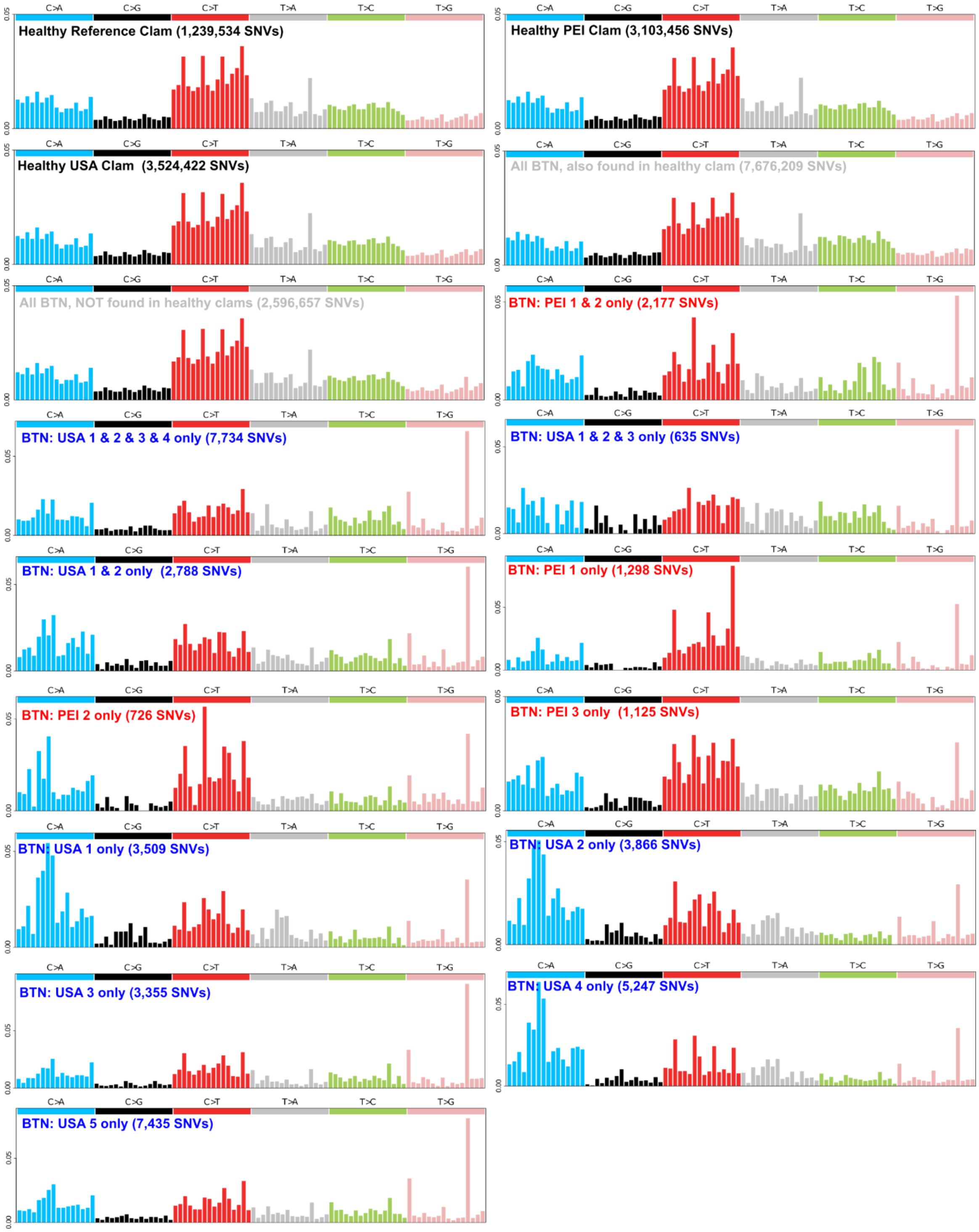


**Supplementary Figure 4: Mutational spectra of SNV from healthy and BTN samples**

Plots show the mutational probability of SNVs in all trinucleotide contexts that were identified in various samples after filtering. Trinucleotide order is the same as shown in Figure 2. Healthy clam SNVs (black labels - top) refer to SNVs that were unique to that clam and not found in other clams, resulting in no overlap of SNVs but still very similar spectra. SNVs found in all BTN samples (grey labels – upper middle) are divided into those found in a healthy clam (likely all from the founder clam genome) and those not found in any of the three healthy clams (includes a mixture of founder and early somatic mutations). Likely somatic SNVs found within the USA (blue labels) and PEI (red labels) sub-lineages show those SNVs that are either shared between all samples (Figure 2b - not shown here), multiple samples (lower middle), or unique to individual samples (bottom). SNVs found in All mutational probabilities are corrected for mutational opportunities in the clam genome, and total mutation counts in each image are shown in the label.

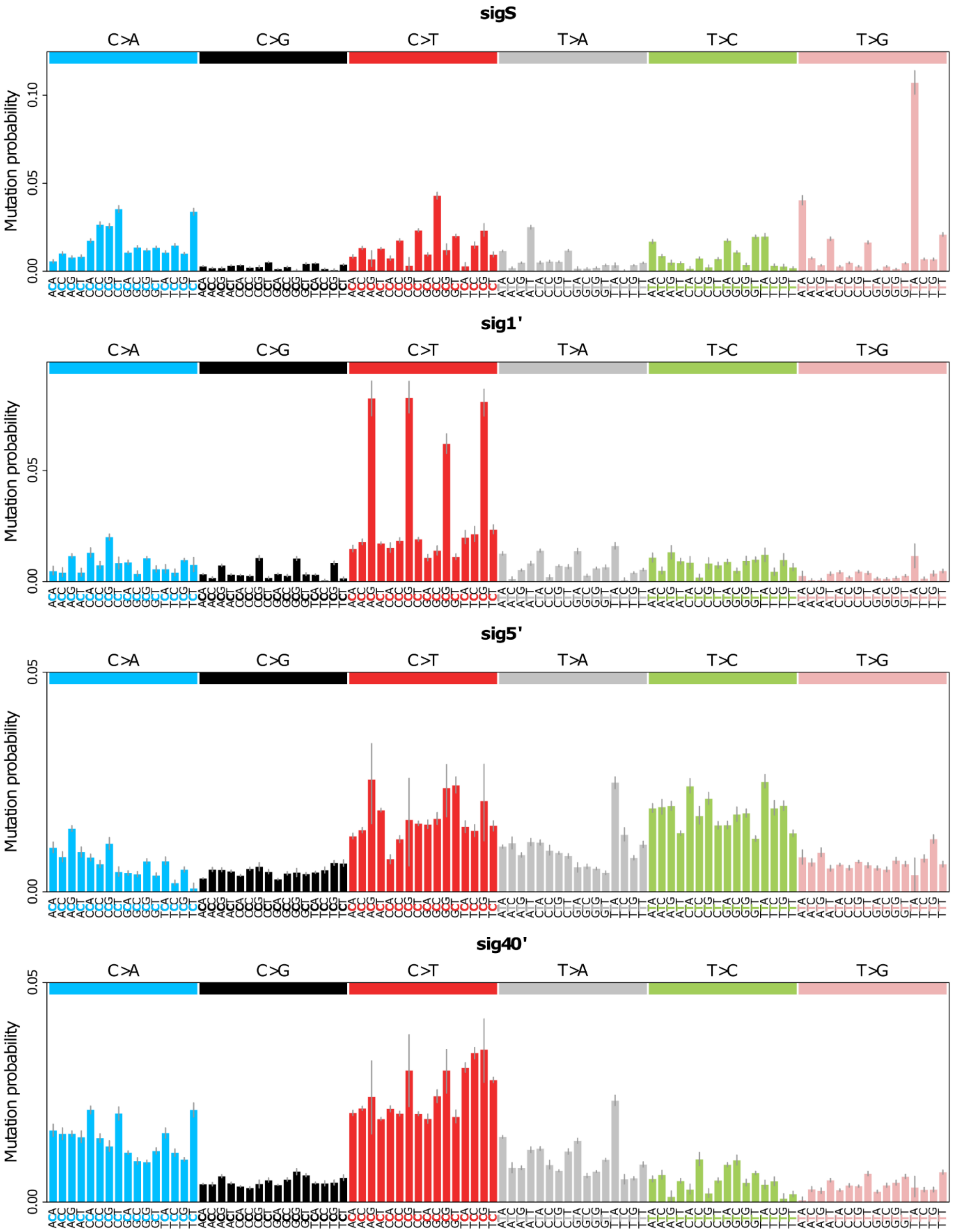


**Supplementary Figure 5: Four mutational signatures identified by *de novo* signature extraction**

We performed *de novo* mutational signature extraction to identify trinucleotide SNV differences between the various samples in this study, yielding four mutational signatures with mutational probabilities corrected for mutational opportunities in the clam genome. Error bars display 95% confidence intervals as determined by the extraction software, sigfit. Signatures sig1', sig5' and sig40' are named after the closest signature in the COSMIC database, as determined by cosine similarity. SigS was named to reflect that it was specific to <u>S</u>omatic mutations in cancer samples.

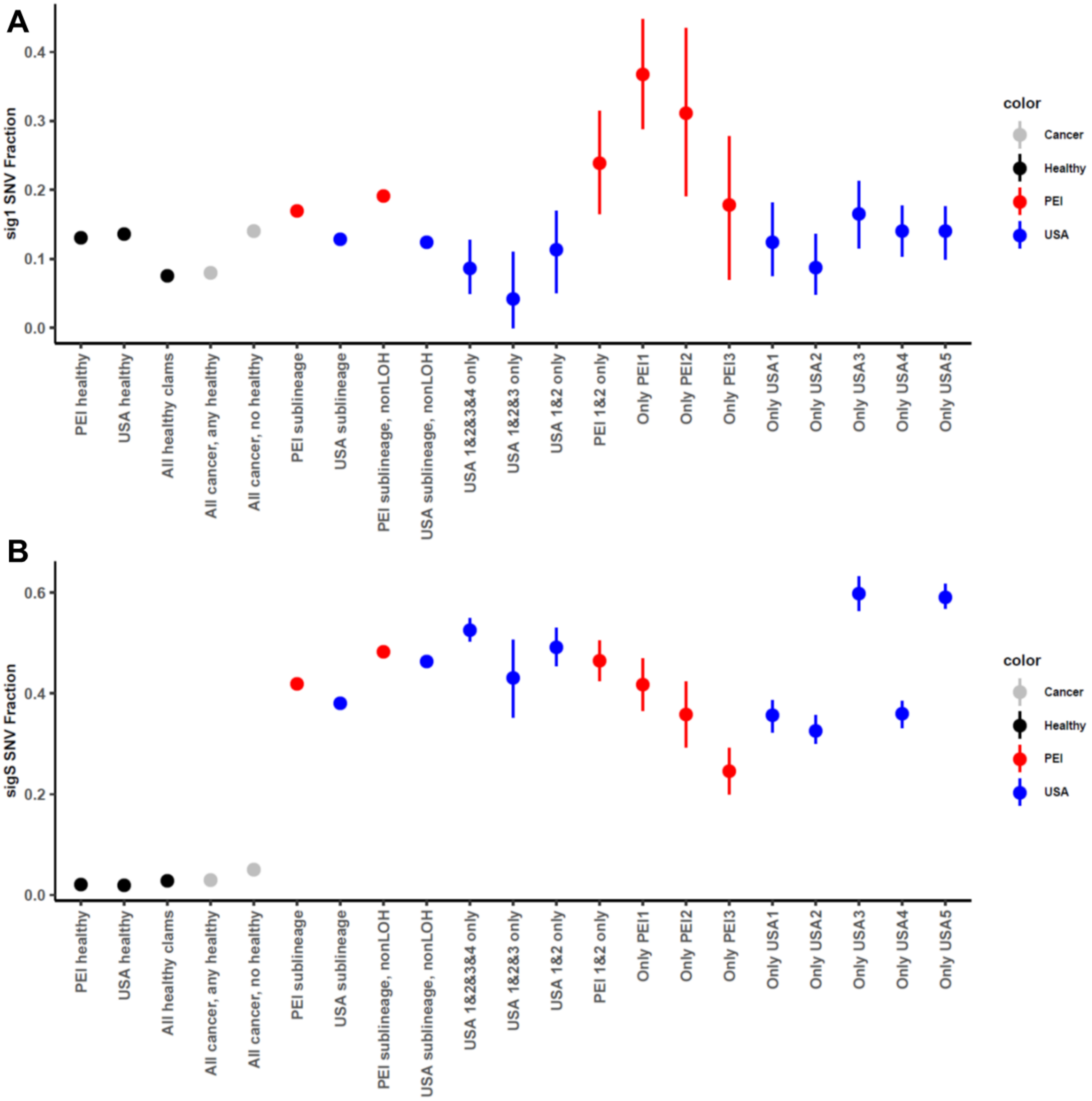


**Supplementary Figure 6: SigS is a large fraction of both USA and PEI, but Sig1 is more highly represented in PEI**
Plots shows the fraction of SNVs attributed to **(A)** signature 1' and **(B)** signature S across healthy and cancer samples, divided and filtered as described in Supplementary Figure 4 and methods (mutational signature extraction and fitting) and diagramed in Supplementary Figure 20. "All healthy clams" refers to SNVs found in all 3 healthy clams in our data set, but not in the reference genome. Error bars display 95% confidence intervals of mutation fractions from fitting error of SNVs to the four mutational signatures.

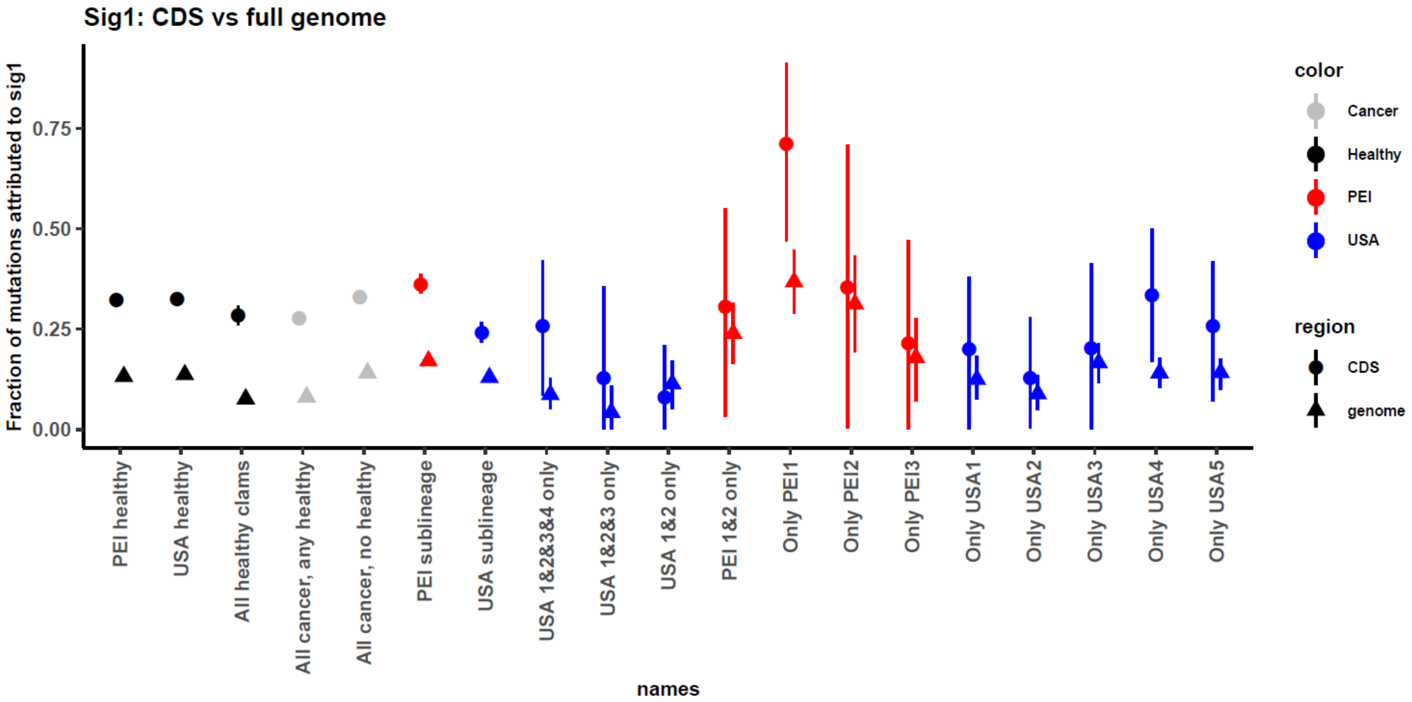


**Supplementary Figure 7: Sig1 is more highly represented in coding regions**

Fraction of mutations attributed to signature 1 across the whole genome (triangles, same data as in supplementary figure 6A) is shown compared to the fraction of signature 1 in coding regions alone (CDS, circles). Note that trinucleotide contexts of mutational opportunities are different in coding regions versus the full genome, which was factored into in the signature fitting process. Error bars display 95% confidence intervals of mutation fractions from fitting error of SNVs to the four mutational signatures.

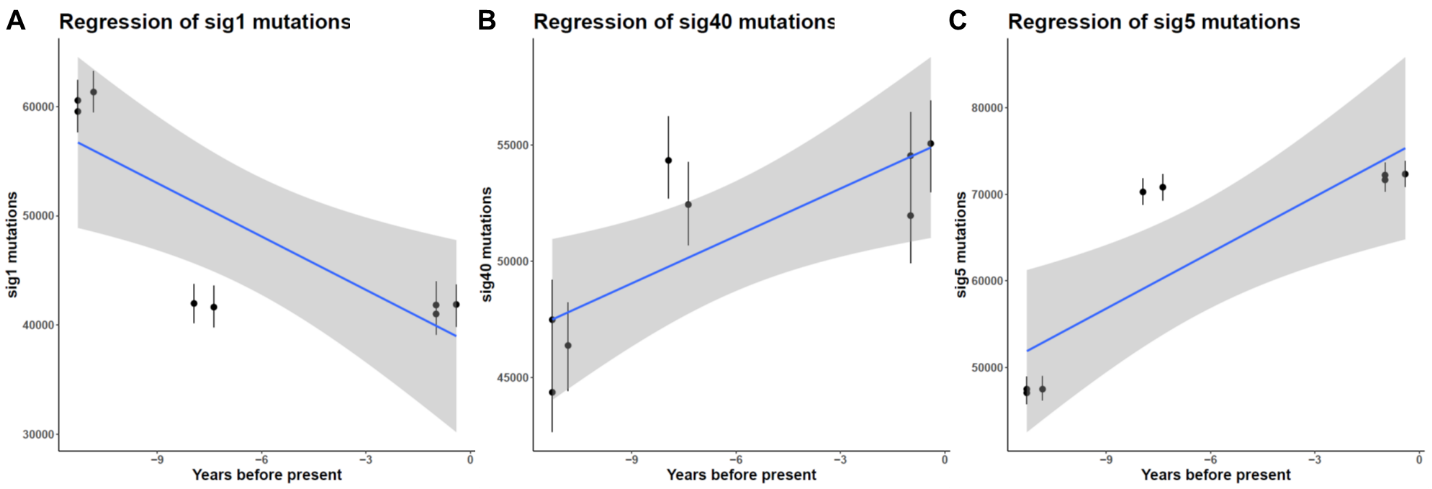


**Supplementary Figure 8: Other signatures do not correlate well with time**

Mutations across MarBTN samples for the other three signatures versus sampling date (SigS is shown in Figure 2C). SNVs found in healthy clams, all BTN samples, or LOH regions are excluded. Note that the earliest three samples (~11 years ago) represent the PEI samples, while the later five samples are USA samples. It is also apparent that sig1'mutation counts are higher in PEI, while sig40' and sig5' mutations are higher in USA. This non-independence of samples leads to trends that are based more on when samples from each subgroup were harvested than mutation accumulation over time. In the case of sigS (as shown in Figure 2C), the correlation holds even when correcting for the relatedness of the samples or when only looking at the USA sub-lineage samples (see methods: cancer dating).

.

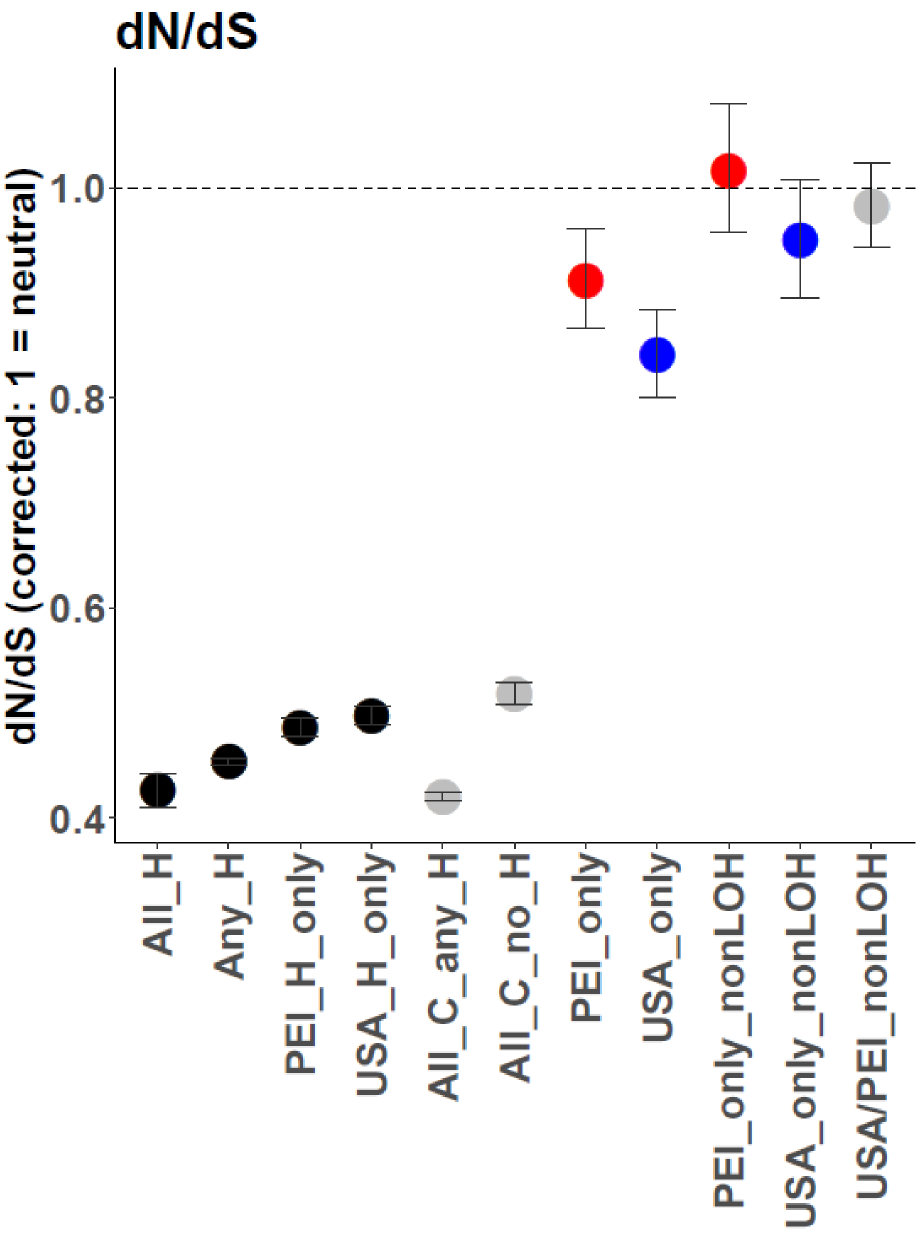


**Supplementary Figure 9: Global dN/dS across germline and somatic SNVs**

dN/dS ratios, corrected so that a ratio of 1 indicates neutrality, across sample bins. Error bars indicate 95% confidence intervals as estimated by dndscv. Sample labels along x-axis are as follows: All_H – SNVs found in all three healthy clams; Any_H – SNVs found in any healthy clam; PEI and USA_H_only – SNVs found in only that healthy clam; All_C_any_H – SNVs found in all cancer samples and at least one healthy clam; All_C_no_H – SNVs found in all cancer samples and no healthy clams; PEI/USA_only – Somatic mutations before excluding LOH regions; PEI and USA_only_nonLOH – High confidence somatic mutations outside putative LOH regions; and USA/PEI_nonLOH – Combined high confidence somatic mutations from both sub-lineages and outside putative LOH regions. After removing LOH regions, dN/dS for high confidence somatic mutations is not significantly different than 1.

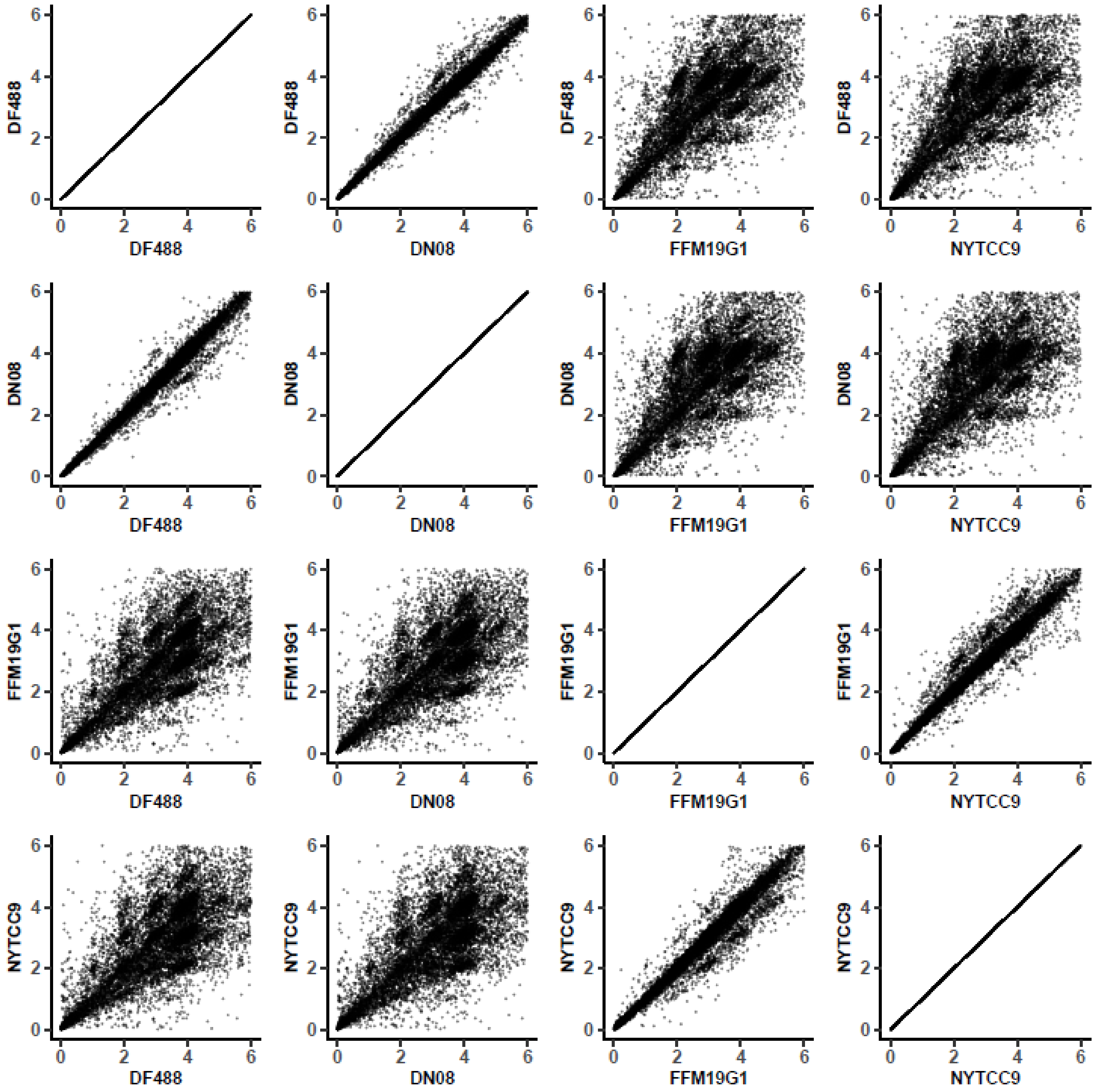


**Supplementary Figure 10: Copy number calls agree closely within sub-lineages, but differ between sub-lineages**
We called copy number across the genome in 100 kB chunks for each sample individually. Here we plot pairwise comparisons of the copy number call for each 100 kB chunk between two representative PEI BTN samples (DN08 and DF488) and two representative USA BTN samples (FFM19G1 and NYTCC9: notably, the two most distantly related USA samples). There is a close correlation ($R^2$>0.94) within sub-lineages (DN08 vs DF488, FFM19G1 vs NYTCC9) and a weaker correlation ($R^2$=0.53-0.56) when comparing between sub-lineages (DN08 or DF488 vs FFM19G1 or NYTCC9). Copy number differences between samples can be seen here as denser groupings of points around integer values that deviate from equal values along the diagonal.

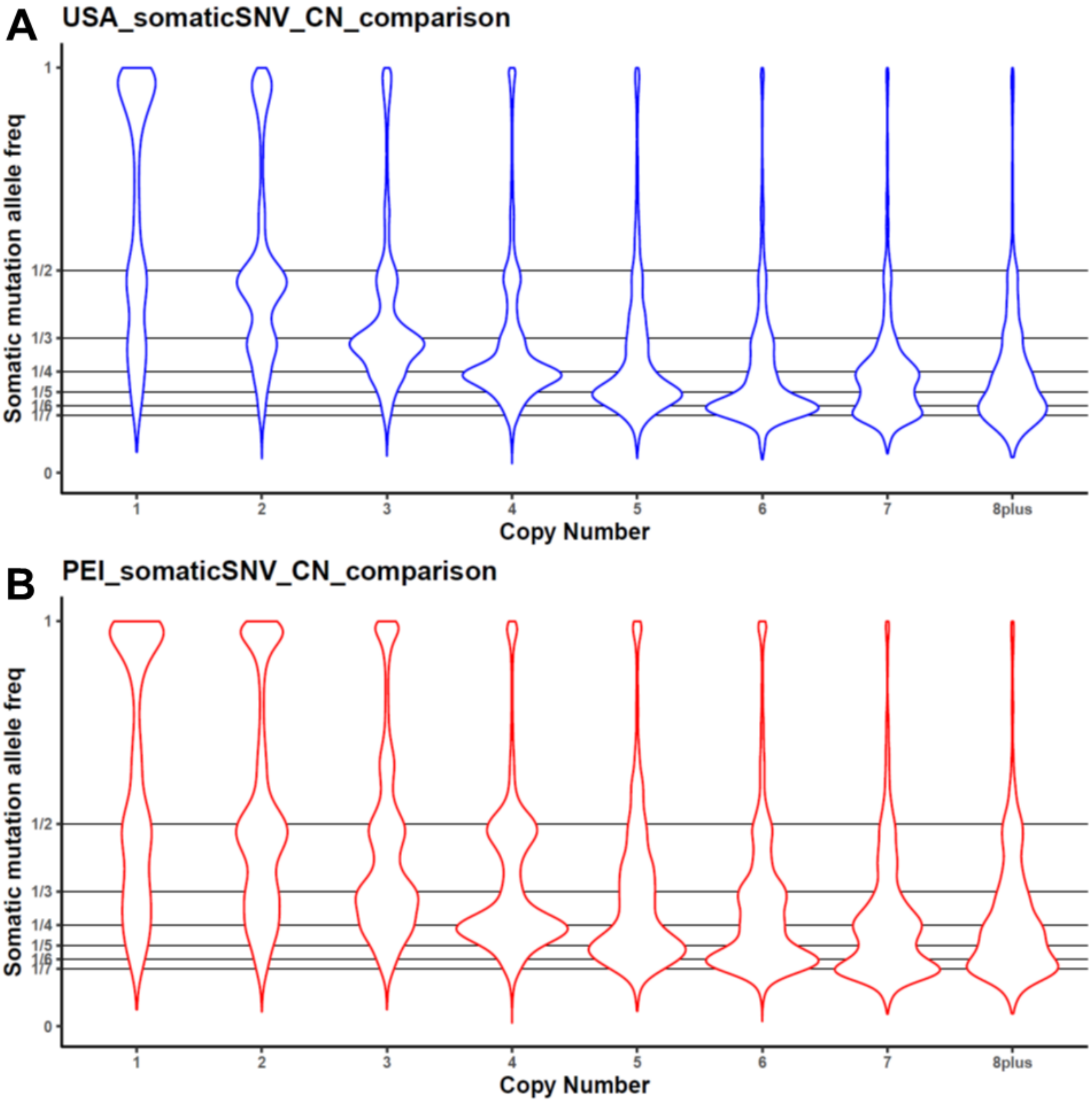


**Supplementary Figure 11: Somatic mutation allele frequencies support copy number calls**

Copy number was called across the genome and the variant allele frequencies of all high confidence somatic mutations were calculated separately for BTN from **(A)** USA) and **(B)** PEI. Violin plots show probability densities of allele frequencies of high confidence somatic mutations, divided into portions of the genome called at each copy number. The peak allele frequency in each case is distributed around the expected value of 1/copy number. In addition to the main, expected peaks for each copy number, in some cases, additional peaks can be seen that indicate somatic mutations prior to copy number gain (e.g. VAF of 0.5 in regions with CN4 that could be due to mutation followed by duplication of the region). Some minor peaks also indicate possible errors in copy number calling or allele frequency counting (e,g, VAF of 0.5 in CN3 regions). These errors could be due to lower read mapping due in polymorphic region, errors caused by repeat regions, regions spanning a CN breakpoint, among other possibilities.

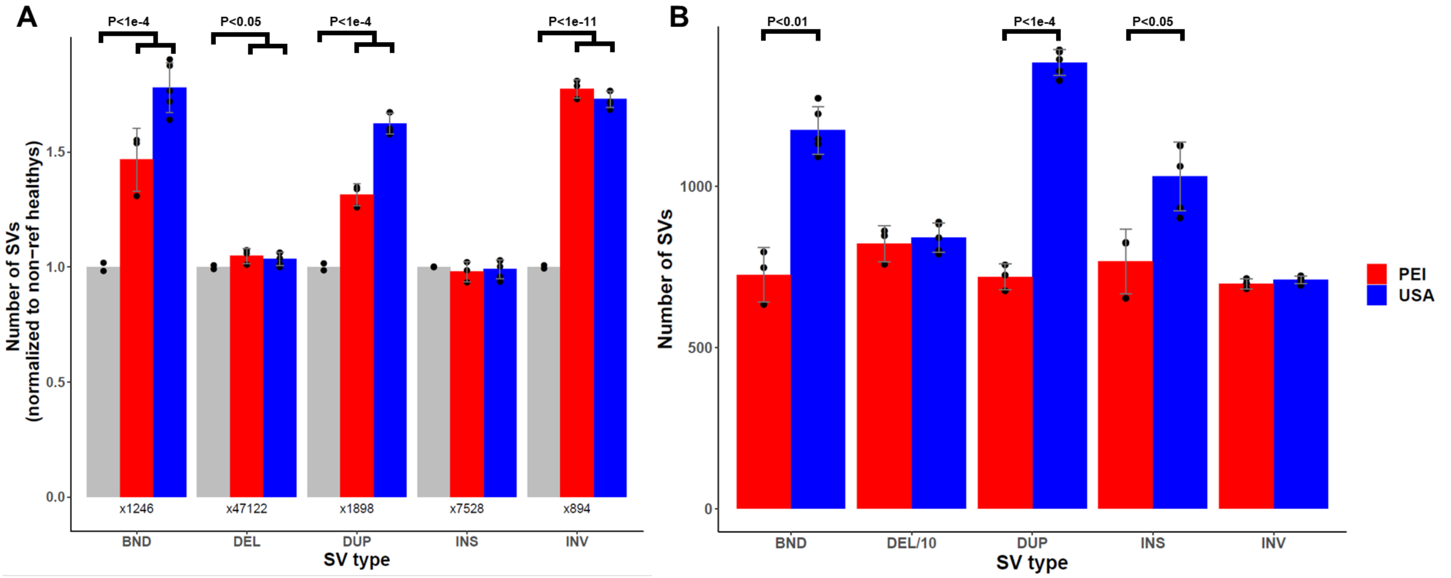


**Supplementary Figure 12: Evidence for elevated somatic structural variants of several types**

Structural variants were called in healthy and BTN samples using Delly, only including only those with precise breakpoints and excluding SVs found in the reference clam. **(A)** The number of called SVs of each type are normalized to the average number of SVs in non-reference healthy clams for each SV type (value above x-axis). Dots represent individual samples, while bars summarize averages for each group: healthy clams, PEI BTN, and USA BTN. Error bars indicate standard deviation. P-values are from unequal variance t-test between BTN samples (n=8) and non-reference healthy clams (n=2). **(B)** Number of called SVs of each type that are unique to each sub-lineage were calculated by removing SVs found in any healthy clams or in any BTN samples from the other sub-lineage. P-values are from unpaired unequal variance t-test between PEI BTN samples (n=3) and USA BTN samples (n=5). Labels follow delly abbreviations of SV types: BND = translocations, DEL = deletions, DUP = tandem duplications, INS = small insertions, INV = Inversions. Deletion counts were much higher than other SV types, so were divided by 10 in (B) for visualization (“DEL/10”).

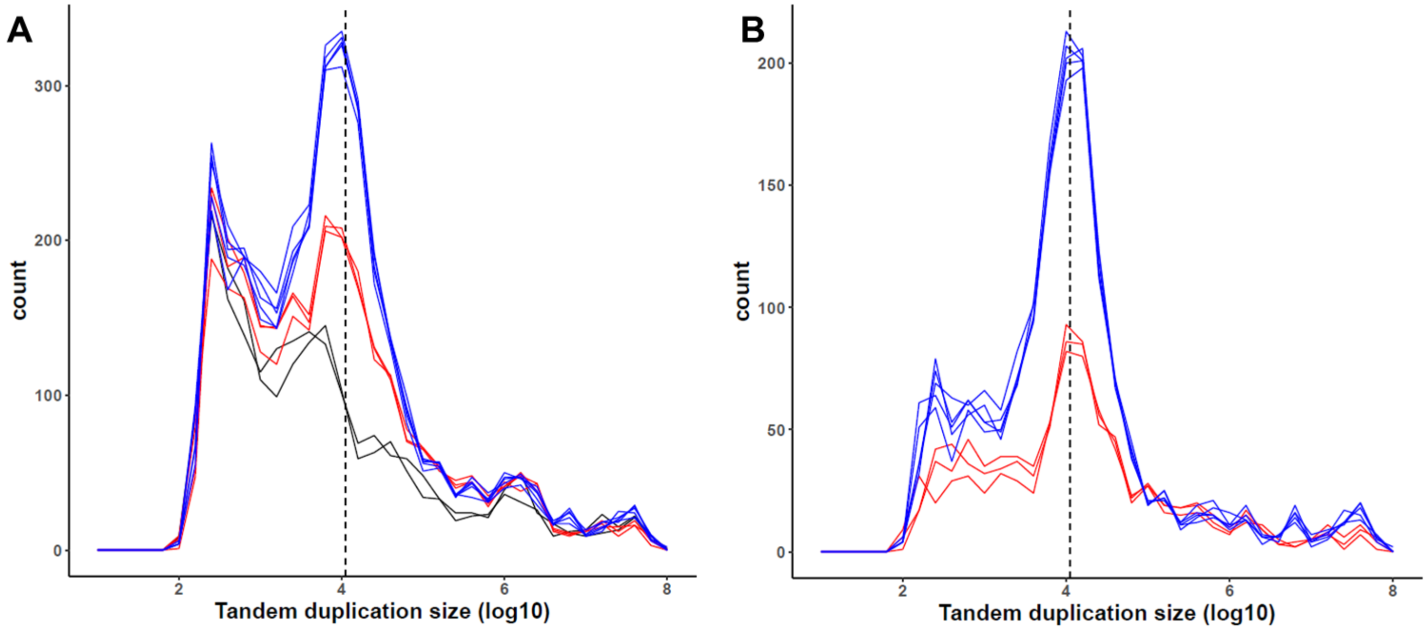


**Supplementary Figure 13: Somatic tandem duplications are distributed around 11 kB**

Plot shows the size distribution of tandem duplications in each sample, after removing SVs found in the reference clam **(A)**, and after removing SVs found in any healthy clams or in any BTN samples from the other sub-lineage **(B)**. Black lines indicate non-reference healthy clams, red lines indicate PEI BTN samples, and blue lines indicate USA BTN samples. Dashed line indicates 11 kB, the median tandem duplication size reported in a tandem duplication phenotype observed in human and mouse cancers with mutant p53 and BRCA1. We observe a bias towards an increase in similarly sized somatic tandem duplications in both sub-lineages of BTN.

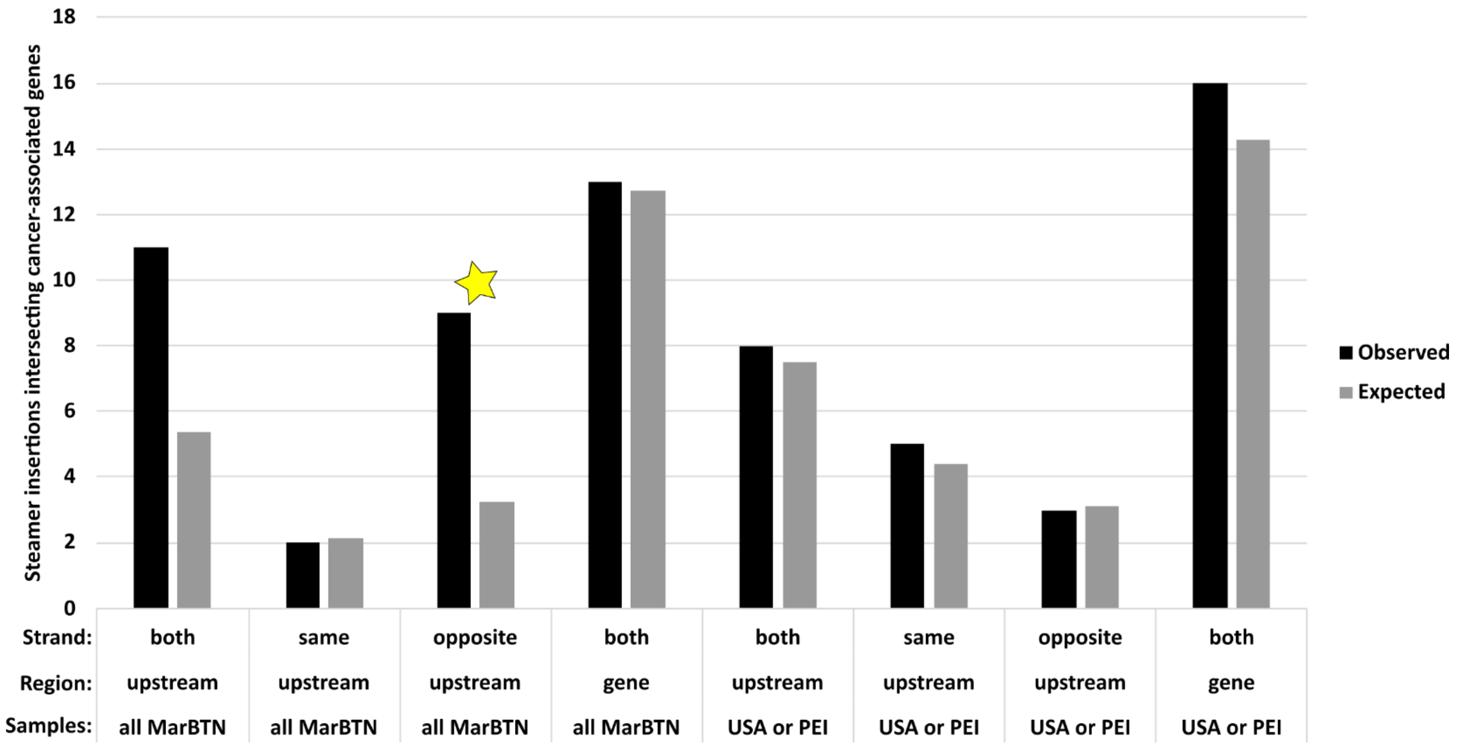


**Supplementary Figure 14: The *Steamer* retrotransposon inserts upstream of cancer-associated orthologs in the opposite direction more often than expected**
We conducted a BLASTP search for the 729 cancer-associated genes in the COSMIC database and found hits in 5,430 of the 38,609 predicted *M. arenaria* genes (14%). If there is not selection for insertion near these genes, we would expect 14% of *Steamer* insertions with a *M. arenaria* gene to intersect with these genes. We counted the number of steamer insertions in genes ("gene") and in the 2 kB upstream genes ("upstream") for early steamer insertions in the lineage trunk ("all MarBTN") and after the divergence of the sub-lineages ("USA or PEI"). We plotted these counts (black) against that expected by chance (grey). Counts match expected closely for late insertions (in only the USA or PEI sub-lineage – right side of plot), either upstream genes or within them, but were higher than expected for early insertions. We further divided upstream insertions by whether the steamer insertion was in the same strand/direction as the gene or opposite, to compare with counts regardless of directionality ("both"). The early insertion bias to insert upstream cosmic genes can be fully explained by a bias to insert in the opposite strand (red star), here with 9/23 (39%) of the genes being cancer associated (would expect 3/23: Chi-squared Bonferroni-corrected p-value = 0.004)

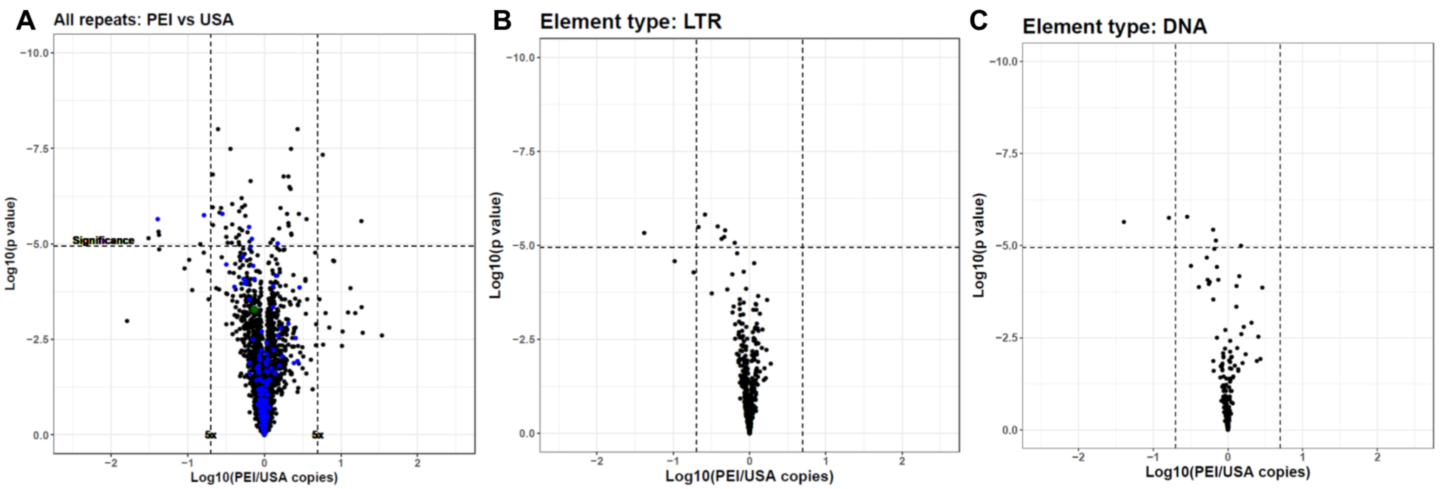


**Supplementary Figure 15: More TEs are expanded in USA vs PEI, particularly LTR elements**

Volcano plot shows estimated copy number of each TE, comparing copy number in MarBTN from PEI with USA for all TE types **(A),** LTR elements **(B**), and DNA transposons **(C)**. TEs more highly amplified in PEI MarBTN are to the right and TEs amplified more highly in USA MarBTN are to the left. Dashed lines correspond to significance threshold (p=0.05, Bonferroni corrected) and 5-fold differences. **(A)** DNA transposons are labeled in blue and *Steamer* is labeled in green. Eight LTR retrotransposons and five DNA transposons are significantly amplified in the USA sub-lineage compared to the PEI sub-lineage, while no identified LTR retrotransposons and a single DNA transposon TEs are significantly amplified in the PEI sub-lineage compared to the USA sub-lineage.

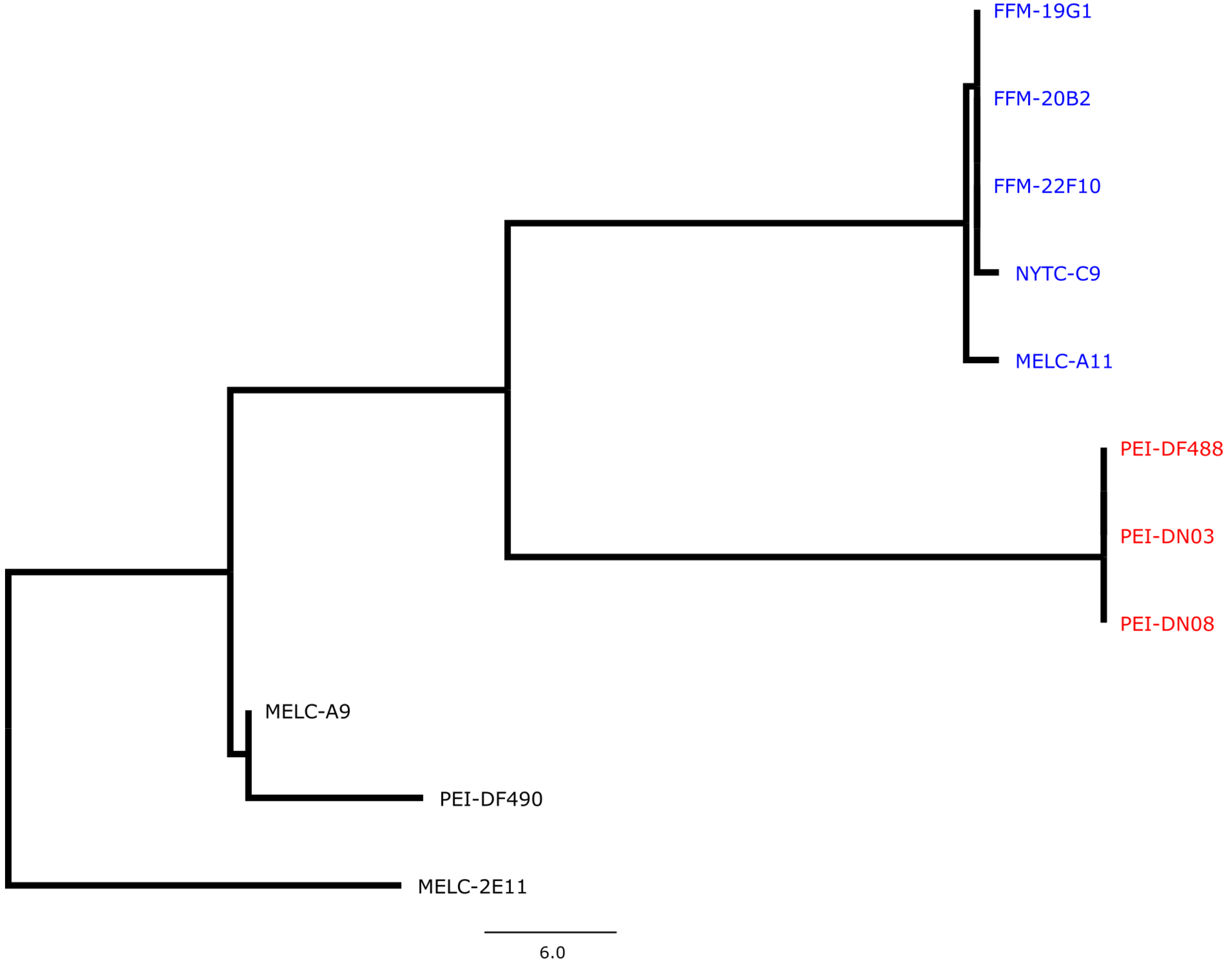


**Supplementary Figure 16: No evidence for mitochondrial transfer**

Phylogenetic tree built from pairwise SNVs called against the previously published *M. arenaria* reference mitogenome (excluding the repeated region) for USA MarBTN samples (blue), PEI MarBTN samples (red) and healthy clams (black). The phylogenetic relationship generally reflects that built from genomic SNVs (i.e., monophyletic MarBTN group with separate USA and PEI sub-lineages).

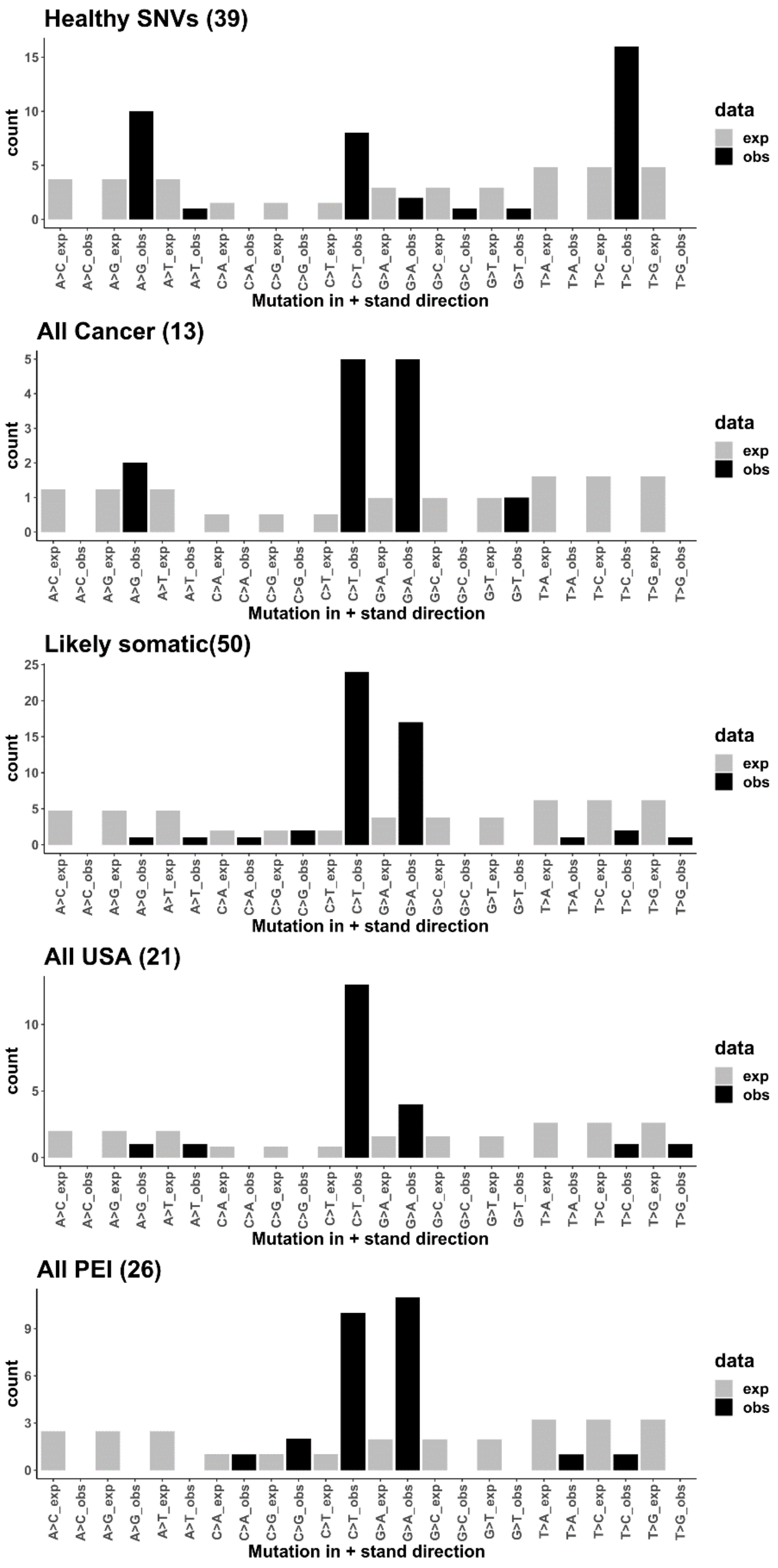


**Supplementary Figure 17: Mitochondria are enriched for transitions in healthy clams, and C>T specifically in somatic mutations**

Observed SNVs (black) compared with expected counts estimated from nucleotide frequencies of the *M. arenaria* mitogenome and assuming equal mutation probability. This calculation was not collapsed to the usual 6 mutation types due to the imbalance of nucleotides in mitochondrial genomes (unequal frequencies of G/C and A/T). Likely somatic refers to SNVs found in a subset of BTN samples, while All USA and All PEI refer to SNVs found in all individuals from that sub-lineage, but not the other sub-lineage.

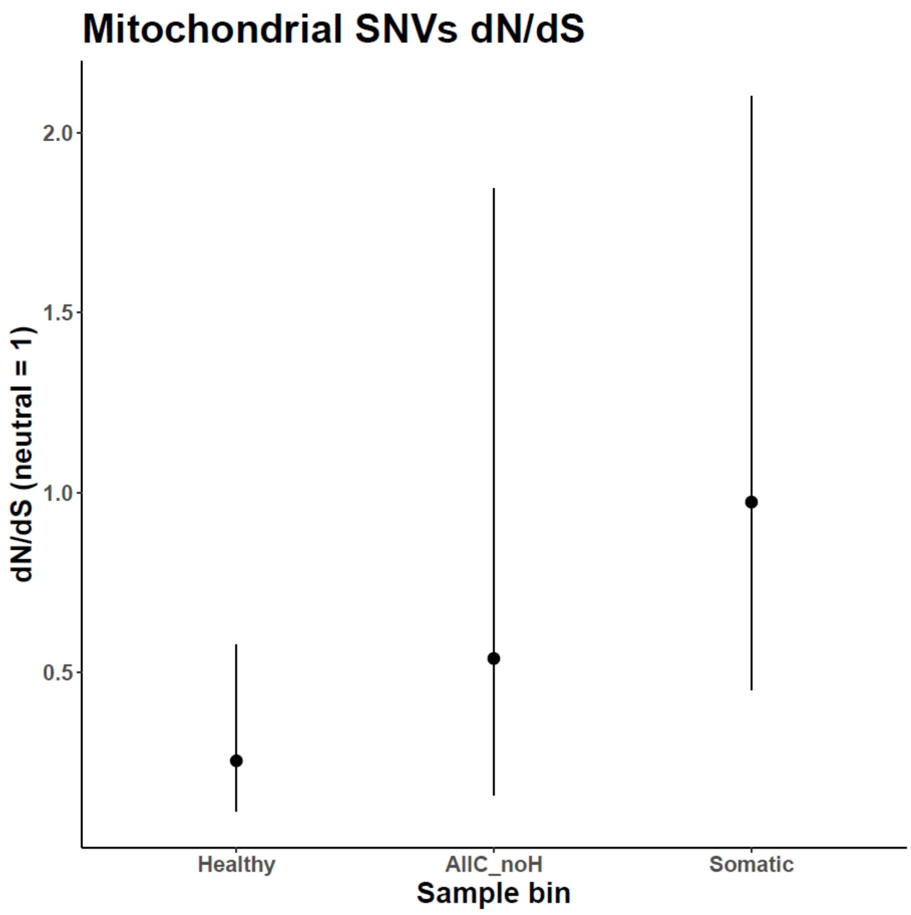


**Supplementary Figure 18: Mitochondria are under relaxed selection in BTN**

dN/dS ratios, where a ratio of 1 indicates neutrality, were calculated for mitochondrial SNVs found in healthy clams, all BTN samples but not healthy clams, and likely somatic mutations (those found in a subset of BTN samples). Error bars indicate 95% confidence intervals as estimated by dndscv and are quite large, due to the low number of mitochondrial SNVs.

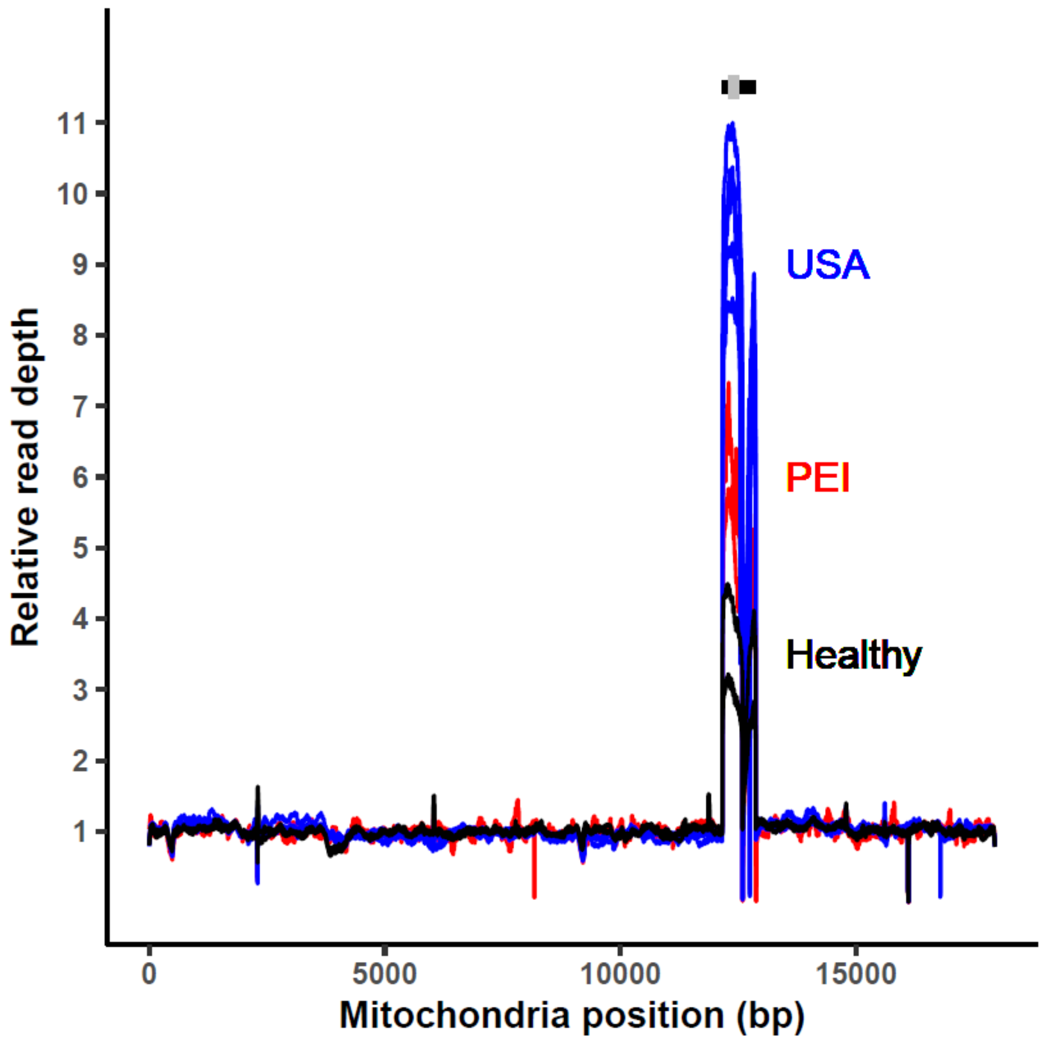


**Supplementary Figure 19. Somatic tandem duplications in mitochondrial D-loop**

Read depth across the mitochondrial genome for healthy clams (black), PEI MarBTN (red) and USA MarBTN (blue), normalized to mean depth outside D-loop. Bars above indicate the D-loop region (12,164-12,870 bp, black) and the region used to estimate duplicated region copy number (12,300-12,500 bp, grey), as shown in Fig 3F.

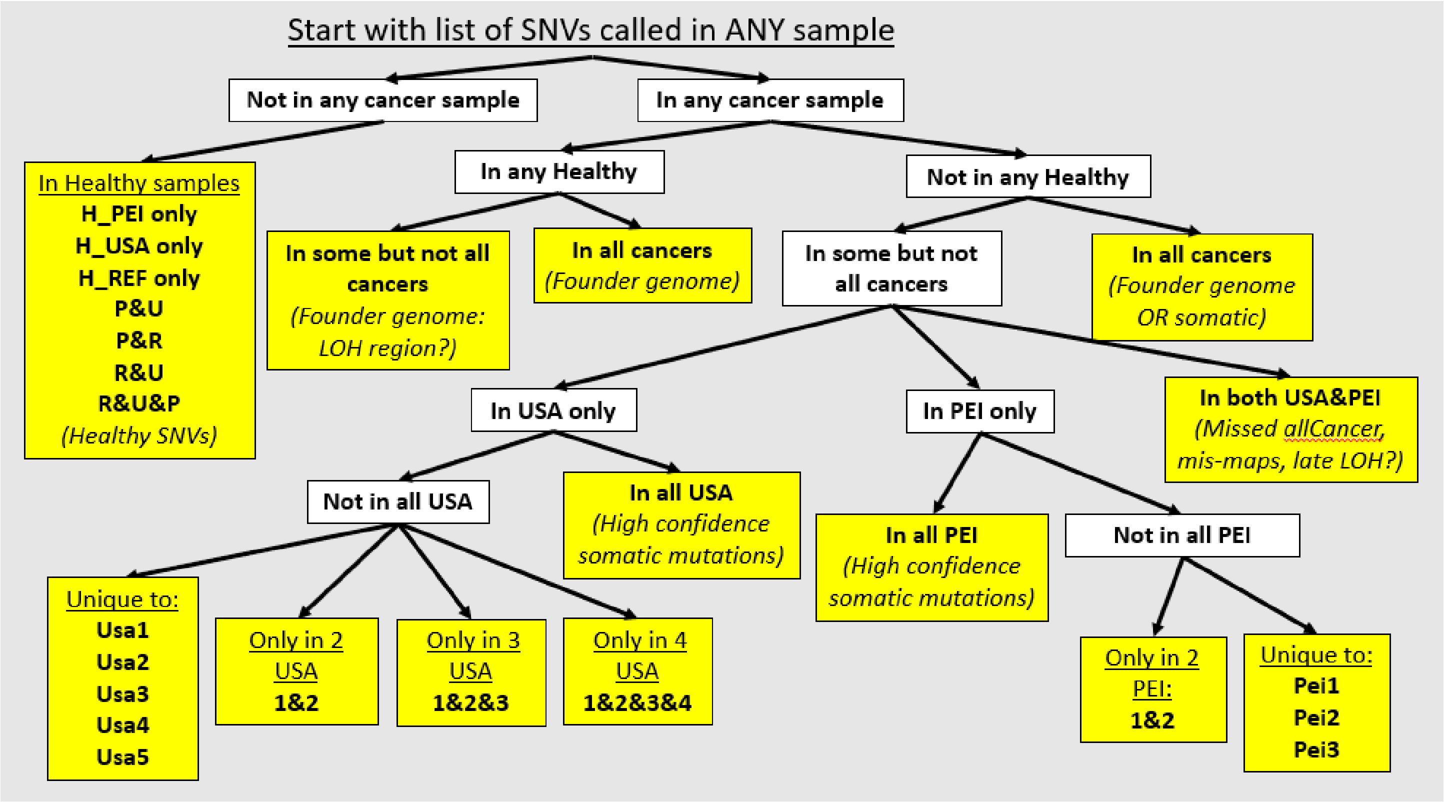


**Supplementary Figure 20: SNV binning strategy for *de novo* signature extraction**

Flowchart of our strategy to separate SNVs into bins for *de novo* signature extraction, based on which sample(s) each SNV was called in. Many of these bins were also used in other analyses, as indicated in the manuscript. The starting point refers to a vcf file of every SNV that was called in at least one of the eleven sample (three healthy, eight cancer) sequenced in this study. Bins highlighted in yellow indicate non-overlapping SNV bins used to for signature extraction.

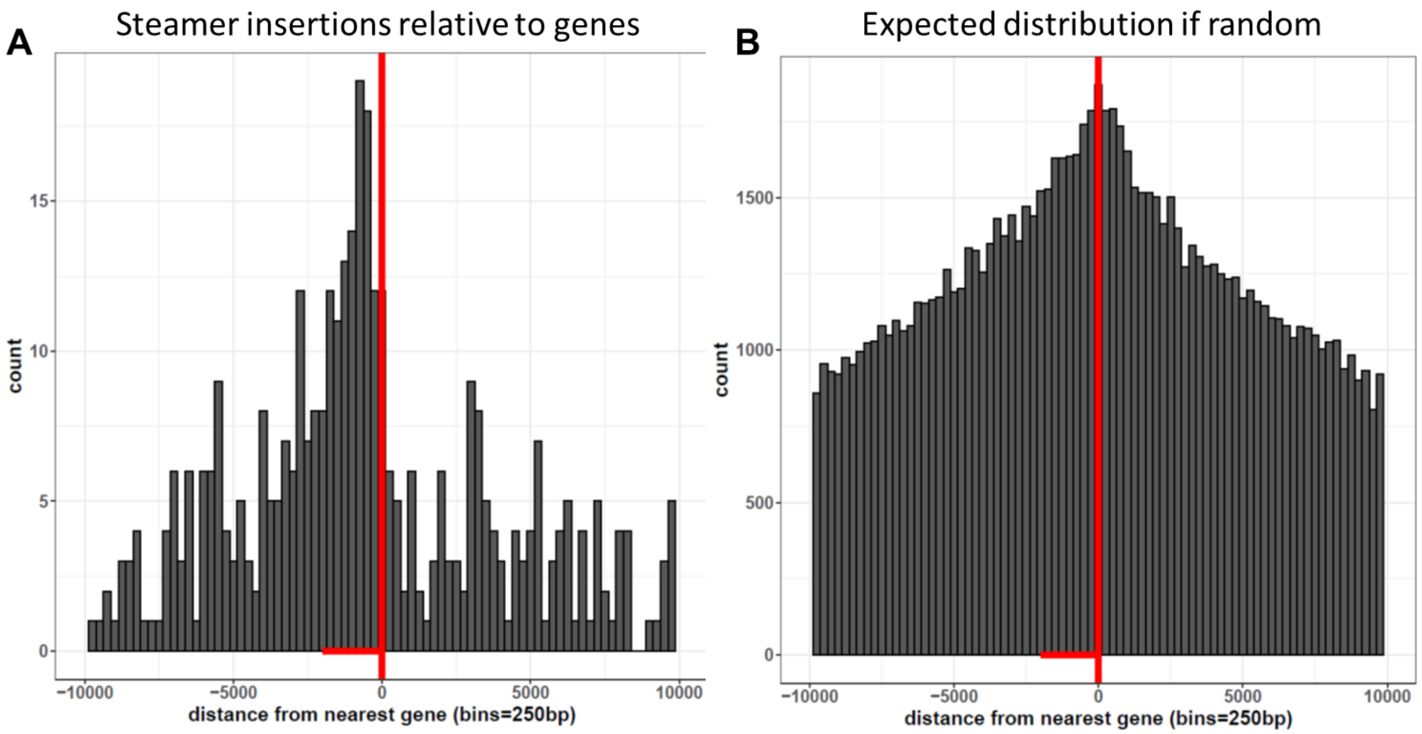


**Supplementary Figure 21: *Steamer* preferentially inserts upstream genes**

**(A)** The histogram shows the distance to nearest gene for *Steamer* insertions found in any cancer sample (n=570). If an insertion was within an annotated gene, the distance to the next nearest insertion was used. 0 (vertical red line) corresponds to the first or last nucleotide of the annotated gene for when the insertion is upstream (negative) or downstream (positive) relative to the gene, respectively. Horizontal red segment highlights 2 kB upstream genes with elevated *Steamer* insertions. **(B)** The histogram shows a distribution of randomly generated insertion sites (n=224,134) based off the observed read mapping in the genome assuming insertions are random.

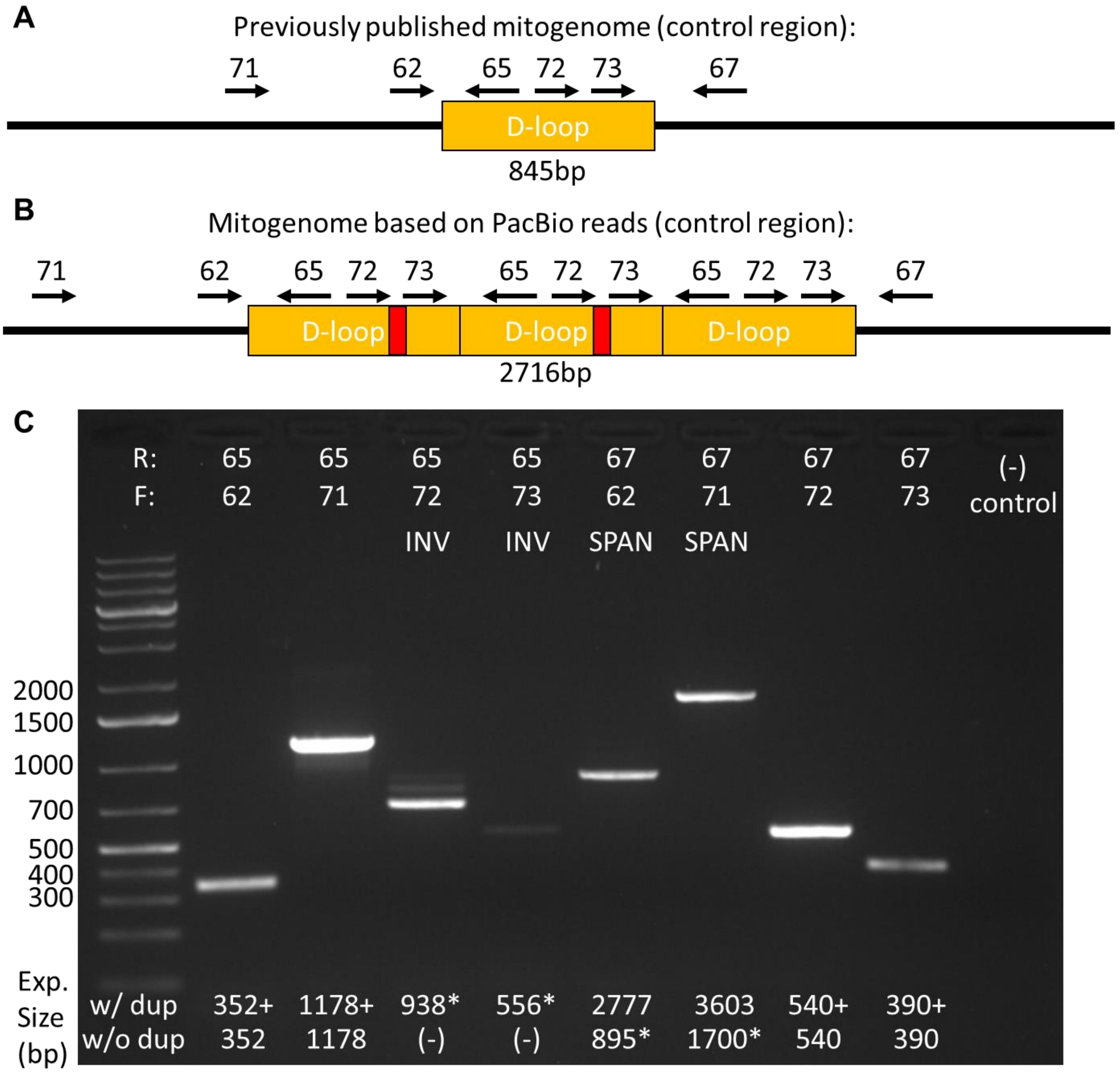


**Supplementary Figure 22: PCR validation of mitochondrial repeat in health clam**

**(A)** Schematic of the control region of the *M. arenaria* control region in the previously published mitogenome with a single d-loop copy and placement of primers (not to scale). **(B)** Schematic of the proposed mitochondrial genome with three d-loop copies and G-rich insertions and placement of primers. **(C)** PCR results. Primer pair combinations are listed on top and expected sizes are listed on bottom. Amplicon sizes from primers spanning the D-loop (67 with 62/71) support a single copy of the D-loop. However, we suspect this is a result of recombination and selection for the smaller product and loss of the G-rich insertions. Inverse PCR with outward-facing primers (65 with 72/72) indicates a tandem duplication allowing outward-facing primers to amplify. The inverse primers spanning the G-rich insertion (65 with 72) has a dim band at expected size, but two brighter bands at smaller sizes.

### SUPPLEMENTAL TABLES

#### Supplementary Table 1: Improved genome contiguity and completeness

| Genome | Sequencing | Source | Length (Gb) | GC (%) | Repeat (%) | Scaffolds | Contigs | Scaffold N50 (kB) | Contig N50 (kB) | BUSCO score (metazoa) |
|---|---|---|---|---|---|---|---|---|---|---|
| Mya.genome.v1.01 | Illumina, low cov PacBio | 2 clams, Plachetzki et al. | 1.32 | 34.98 | 35.54 | 152,330 | 226,958 | 14.7 | 10.6 | C:71.4%[S:55.6%,D:15.8%],F:16.7%,M:11.9%,n:954 |
| Mar.3.1.1 | 10x | MELC-2E11, This paper | 1.29 | 35.27 | 39.64 | 1,029,422 | 1,100,210 | 22.1 | 11.0 | C:73.9%[S:62.8%,D:11.1%],F:13.1%,M:13.0%,n:954 |
| **Mar.3.4.6.p1** | **PacBio, HiC, 10x** | **MELC-2E11, This paper** | **1.22** | **35.32** | **41.72** | **17** | **539** | **58,023** | **3,381** | **C:94.9%[S:92.5%,D:2.4%],F:1.2%,M:3.9%,n:954** |

**Supplementary Table 2: List of sequenced samples**

| Name | Map code | Alternate aliases | Healthy/BTN | Date sampled | Location |
|---|---|---|---|---|---|
| MELC-2E11 | | | Healthy | 6/1/2018 | Larrabe Cove, Machiasport, ME, USA |
| MELC-A9* | | | Healthy | 9/18/2013 | Larrabe Cove, Machiasport, ME, USA |
| PEI-DF490* | | Dfar490 | Healthy | 5/1/2010 | Dunk Estuary, PEI, Canada |
| PEI-DF488* | pei1 | DF-488, Dfar-488 | BTN | 10/28/2010 | Dunk Estuary, PEI, Canada |
| PEI-DN03* | pei2 | DN-HL03, Dnear-HL03 | BTN | 5/1/2010 | Dunk Estuary, PEI, Canada |
| PEI-DN08* | pei3 | Dnear-08 | BTN | 5/1/2010 | Dunk Estuary, PEI, Canada |
| FFM-19G1 | usa1 | | BTN | 8/31/2020 | Brunswick, ME, USA |
| FFM-20B2 | usa2 | | BTN | 8/31/2020 | Brunswick, ME, USA |
| FFM-22F10 | usa3 | | BTN | 3/31/2021 | Waldoboro, ME, USA |
| MELC-A11* | usa4 | | BTN | 9/18/2013 | Larrabe Cove, Machiasport, ME, USA |
| NYTC-C9* | usa5 | | BTN | 4/12/2014 | Long Island Northshore, NY, USA |

*previously reported in Metzger *et al.* 2015

#### Supplementary Table 3: dN/dS positive selection hits table

| *Gene annotation information* | | | |
|---|---|---|---|
| **annotated_name** | **maker_name** | **AA #** | **top bivalve blastp hit** |
| uncharacterized_1199 | maker-Mar.3.4.6.p1_scaffold15-snap-gene-127.5 | 312 | no hits |
| TEN1-like_3 | maker-Mar.3.4.6.p1_scaffold9-snap-gene-700.19 | 112 | receptor-type tyrosine-protein phosphatase mu-like (Mizuhopecten yessoensis) |
| uncharacterized_7181 | maker-Mar.3.4.6.p1_scaffold11-snap-gene-477.12 | 80 | no hits |
| uncharacterized_10146 | maker-Mar.3.4.6.p1_scaffold4-snap-gene-785.0 | 211 | uncharacterized protein (Crassostrea virginica) |
| PIF1-like_47 | maker-Mar.3.4.6.p1_scaffold6-snap-gene-524.0 | 447 | ATP-dependent DNA helicase PIF1 (Mytilus galloprovincialis) |

| | *Somatic mutations* | | | | *Healthy clam SNVs* | | | |
|---|---|---|---|---|---|---|---|---|
| **annotated_name** | **n_syn** | **n_mis** | **wmis_cv** | **qmis_cv** | **n_syn** | **n_mis** | **wmis_cv** | **qmis_cv** |
| uncharacterized_1199 | 2 | 30 | 35.963162 | 2.42E-08 | 0 | 1 | 0.05840293 | 9.22E-07 |
| TEN1-like_3 | 2 | 12 | 34.868098 | 1.64E-04 | 13 | 14 | 0.42658891 | 4.53E-02 |
| uncharacterized_7181 | 1 | 9 | 105.558072 | 1.64E-04 | 3 | 3 | 0.57100099 | 5.11E-01 |
| uncharacterized_10146 | 4 | 15 | 11.320769 | 9.25E-03 | 4 | 0 | 0 | 1.06E-05 |
| PIF1-like_47 | 17 | 41 | 3.491338 | 2.42E-02 | 10 | 35 | 1.33074384 | 4.77E-01 |

AA #: gene length in amino acids
n_syn: number of synonomous SNVs
n_mis: number of misense SNVs
wmis_cv: dN/dS ratio after corrections
qmis_cv: significance after corrections

**Supplementary Table 4: Primers for inverse PCR**

| ID code | Oligo name | Sequence | Direction |
|---|---|---|---|
| SHO_062 | SHO_062_MyaMT_dloop_F1 | TACGAGCAAAAGCCGTTCCT | F |
| SHO_065 | SHO_065_MyaMT_dloop_R1 | CCCATAACGCCCGATTTTGC | R |
| SHO_067 | SHO_067_MyaMT_dloop_R3 | AACCGAGCTGACCTCATTCA | R |
| SHO_071 | SHO_071_MyaMT_dloop_F7 | TCCTGTGTGCCGAAAGAGTC | F |
| SHO_072 | SHO_072_MyaMT_dloop_F8 | CGTGGCGGGAGTATACAGTG | F |
| SHO_073 | SHO_073_MyaMT_dloop_F9 | GGAGAGGGGAGAGGGGATTT | F |
